# PVRL2 Suppresses Anti-tumor Immunity Through PVRIG- and TIGIT-Independent Pathways

**DOI:** 10.1101/2024.01.26.577132

**Authors:** Jiuling Yang, Li Wang, James R. Byrnes, Lisa L. Kirkemo, Hannah Driks, Cassandra D. Belair, Oscar A. Aguilar, Lewis L. Lanier, James A. Wells, Lawrence Fong, Robert Blelloch

**Affiliations:** Department of Urology, University of California San Francisco, San Francisco, CA 94143 USA; Department of Pharmaceutical Chemistry, University of California San Francisco, San Francisco, CA 94158 USA; Department of Microbiology and Immunology, University of California, San Francisco, and Parker Institute for Cancer Immunotherapy, San Francisco, CA 94143, USA; Division of Hematology/Oncology, Department of Medicine, University of California, San Francisco, San Francisco, CA 94143 USA

**Author notes:** **Corresponding Author**: Robert Blelloch, University of California San Francisco, 35 Medical Center Way, RMB1018, San Francisco, California, 94143.

## Abstract

PVRL2 is believed to act as an immune checkpoint protein in cancer; however, most insight into PVRL2’s role is inferred from studies on its known receptor PVRIG. Here, we directly study PVRL2. PVRL2 levels are high in tumor cells and tumor-derived exosomes. Deletion of PVRL2 in multiple syngeneic mouse models of cancer shows a dramatic reduction in tumor growth that is immune dependent. This effect can be even greater than seen with deletion of PD-L1. PVRL2 functions by suppressing CD8 T and NK cells in the tumor microenvironment. Unexpectedly, the effect of PVRL2 loss on tumor growth remains in the absence of PVRIG. In contrast, PVRIG loss shows no additive effect in the absence of PVRL2. TIGIT blockade combined with PVRL2 deletion results in the greatest reduction in tumor growth. This effect is not recapitulated by the combined deletion of PVRL2 with its paralog PVR, the ligand for TIGIT. These data uncover PVRL2 as a distinct inhibitor of the anti-tumor immune response with functions beyond that of its known receptor PVRIG. Importantly, the data provide a strong rationale for combinatorial targeting of PVRL2 and TIGIT for cancer immunotherapy.

## Introduction

Over the past decade, immune checkpoint inhibitors (ICIs), including antibodies blocking immune checkpoints PD-1, PD-L1, and CTLA-4, have made significant progress in advancing cancer immunotherapy. These ICIs enhance the host immune system to combat cancer and have achieved remarkable success across numerous cancer types. Nevertheless, only 10-30% of cancer patients exhibit favorable responses to these therapies (1–4). Moreover, a majority of patients who initially respond eventually develop resistance during the course of treatment (1, 4). The underlying mechanisms responsible for the initial or acquired resistance in a large percentage of patients remain mostly unknown. Therefore, it is of utmost importance to gain a better understanding of these mechanisms and identify additional immunotherapeutic strategies to overcome resistance to further improve cancer care.

Exosomes have emerged as one such potential mechanism of resistance (5–7). Exosomes are small extracellular vesicles (EVs) ranging from 50-150 nm in diameter that originate from the endosome system of almost all mammalian cells, including tumors cells (8). Recent studies from our group and several others have demonstrated that tumor-derived exosomes can present PD-L1 suppressing T cell activation and promoting tumor growth across multiple cancer types (5, 6, 9, 10). However, PD-L1 is unlikely to fully explain the immunosuppressive properties of exosomes as anti-PD-L1/PD-1 blockade fails to fully recapitulate the loss of exosomes on the anti-tumor immune response and tumor growth (5, 6). Therefore, there is need to better understand the mechanisms underlying exosome driven immune suppression.

In recent years, there has been a rapidly growing list of potential immune checkpoint proteins including members of the Nectin and Nectin-like family. In particular, the ligands PVRL2 (also known as Nectin-2 or CD112) and PVR (also known as Necl-5 or CD155) have been proposed to play immunoregulatory roles in tumor progression (11, 12). PVRL2 and PVR are expressed on tumor cells, antigen-presenting cells (APCs), and endothelial cells (13). They interact with a co-stimulatory receptor DNAX accessory molecule 1 (DNAM-1, also known as CD226), which is expressed on T cells and NK cells, stimulating their activation (14–16). However, they also bind immunoinhibitory receptors, including the T cell immunoreceptor with Ig and ITIM domains protein (TIGIT), poliovirus receptor–related immunoglobulin domain protein (PVRIG, also called CD112R), and CD96 (also known as TACTILE) (17–22). PVR serves as the primary ligand for TIGIT, while PVRL2 is thought to be the primary ligand for PVRIG (17–19). PVRL2 may also bind to TIGIT, but with low affinity (17, 18). PVR is also a ligand for CD96, although its role in regulating immune cells remains unclear due to conflicting results (20–23). Similar to other checkpoint receptors such as PD-1, both TIGIT and PVRIG contain an immunoreceptor tyrosine-based inhibitory motif (ITIM) within their cytoplasmic tails (14, 17, 18), albeit truncated in mouse PVRIG (24). The binding affinities of PVR to TIGIT and PVRL2 to PVRIG are much higher than their affinities to DNAM-1 (17, 19, 25, 26). Thus, TIGIT and PVRIG may also function in part by out-competing DNAM-1 for ligand binding (26, 27); however, mechanistic studies on these pathways are rather limited.

In recent years, the PVR-TIGIT and PVRL2-PVRIG pathways have been pursued as potential novel therapeutic targets for cancer. Blocking antibodies against TIGIT, PVRIG, and PVR have been developed and are currently in various stages of clinical trials. TIGIT was the first target to be evaluated for therapeutic development and many anti-TIGIT antibodies are currently undergoing phase I-III clinical trials, showing promising outcomes particularly when combined with anti-PD-1 or PD-L1 therapies (28). More recently, an anti-PVRIG antibody, COM701, has entered phase I clinical trials as monotherapy and in combination with an anti-PD-1 antibody (nivolumab) (NCT03667716), and with nivolumab and an anti-TIGIT antibody (BMS-986207) (NCT04570839) for solid tumors (29, 30). Another anti-PVRIG antibody, NM1F also entered phase I clinical trial in 2023 for advanced solid tumors (NCT05746897). Furthermore, an anti-PVR antibody has been developed and entered in phase I clinical trials in 2022 (NCT05378425). No such efforts have been taken for PVRL2, likely at least in part because its functions are thought to be through PVRIG and thus targeting it could be considered redundant with current anti-PVRIG development strategies.

To date, very few studies have directly evaluated the role of PVRL2 in anti-tumor immunity and assessed its potential as a therapeutic target. Using mass spectrometry-based proteomics, we not only identified PVRL2 on tumor cells, but also tumor-derived exosomes. We performed follow-up genetic studies to show a partial role for PVRL2 on exosomes in promoting tumor growth, and an even larger role on cells. Loss of PVRL2 showed a dramatic reduction in tumor growth by impacting both the adaptive and innate immune responses, while PVR mostly impacted the innate immune response. Surprisingly, these effects were largely independent of PVRIG. Combinatorial inhibition of TIGIT, but not PVR, and loss of PVRL2 showed the largest effects. These data uncover new roles for PVRL2 in the anti-tumor immune suppression and provide a strong rationale for targeting PVRL2 as a novel strategy in cancer care.

## Materials and Methods

### Cell lines

Human tumor cell lines: PC3 and SK-MEL-28 cell lines were purchased from ATCC. PC3 is a prostatic adenocarcinoma cell line derived from a male patient (31). SK-MEL-28 is a malignant melanoma cell line isolated from a male patient (32). PC3 cells were cultured in F-12K Medium (Kaighn’s Modification of Ham’s F-12 Medium) (GIBCO, ref. 21127– 022), supplemented with 10% Fetal Bovine Serum (Corning, ref. 35-010-CV) and Penicillin/Streptomycin (Sigma, cat. P4333). SK-MEL-28 cells were cultured in ATCC-formulated Eagle’s Minimum Essential Medium (ATCC, cat. 30-2003) with 10% Fetal Bovine Serum (Corning, ref. 35-010-CV) and Penicillin/Streptomycin (Sigma, cat. P4333). All cells were cultured at 37°C in a humidified atmosphere containing 5% CO_2_.

Mouse tumor cell lines: TRAMP-C2, CT26 and B16F10 cells were obtained from ATCC. MC38 cells were kindly provided by Jeffrey Schlom’s Lab at the National Cancer Institute (NCI) at National Institutes of Health (NIH). TRAMP-C2 cells were transgenic prostate adenocarcinoma cells derived from a C57BL/6 male mouse (33). CT26 cells are undifferentiated murine colon carcinoma cells derived from a female BALB/c mouse induced with N-nitroso-N-methylurethane-(NNMU) (34). B16F10 cells are a melanoma cell subline from B16 parental line derived from a male C57BL/6 mouse that has high lung metastatic ability (35). MC38 are murine colon adenocarcinoma cells derived from a female C57BL/6 (36).

TRAMP-C2 cells were cultured in Dulbecco’s Modified Eagle’s Medium (DMEM) (UCSF cell culture facility), with 5% Nu-Serum IV (Corning, cat. 80089-542), 5% Fetal Bovine Serum (Corning, ref. 35-010-CV), 0.005 mg/mL Bovine Insulin (Sigma, cat. I0516), 10 nM dehydroepiandrosterone (DHEA) (Sigma, cat. D-063), and Penicillin/Streptomycin (Sigma, cat. P4333). B16F10 cells were cultured in ATCC-formulated Dulbecco’s Modified Eagle’s Medium (ATCC, cat. 30-2002) with 10% Fetal Bovine Serum (Corning, ref. 35-010-CV) and Penicillin/Streptomycin (Sigma, cat. P4333). CT26 cells were cultured in RPMI-1640 Medium (GIBCO, ref. A10491-01) with 10% Fetal Bovine Serum (Corning, ref. 35-010-CV) and Penicillin/Streptomycin (Sigma, cat. P4333). MC38 cells were cultured in DME H-21 (Dulbecco’s Modified Eagle Medium) High Glucose (UCSF cell culture facility) containing 10% Fetal Bovine Serum (Corning, ref. 35-010-CV), Penicillin/Streptomycin (Sigma, cat. P4333), 1 mM sodium pyruvate (GIBCO, ref. 11360-070), 1% NEAA (GIBCO, ref. 11140-050), 0.05 mg/mL Gentamicin (GIBCO, ref. 15750-060). All cells were cultured at 37°C in a humidified atmosphere containing 5% CO_2_.

### Mouse strains

WT C57BL/6 mice (Stock # 000664) and Balb/cJ mice (Stock # 000651) were purchased from The Jackson Laboratory. Immunodeficient NOD CRISPR *Prdkc Il2r Gamma* (NCG) mice were purchased from Charles River (Stock # 572). The *Pvrig* KO mice were derived from sperm obtained through the NIH Knockout Mouse Project (KOMP) program from the Mutant Mouse Regional Resource Centers (MMRRC) at UCDavis (Stock # 043995-UCD). *In vitro* fertilization (IVF) was performed at the University of California San Francisco (UCSF) Cryopreservation Core. The resulting mice were bred and genotyped by PCR (primer sequences are listed below). *Rag1* KO mice were kindly shared by Alexander Marson’s lab at UCSF and were originally purchased from The Jackson Laboratory (Stock # 002216). Age matched male mice ranging 8-11 weeks old were used for all experiments. Mice were randomly assigned to experimental or control groups. Mice were bred and housed under specific-pathogen-free condition. All experiments were conducted under the pre-approved protocols by the Institutional Animal Care and Use Committee at UCSF (protocol # AN188927) and guidelines set by the National Institutes of Health (NIH).

*Pvrig* KO genotyping common forward primer: GTTCCATTCCCTGCCCCTTAGC

*Pvrig* KO genotyping common reverse primer: CGTACTCTTCGGCTCACACTTGTGT

*Pvrig* KO genotyping WT reverse primer: GCAATGTTGAGAATAGAACCAGGGTC

### Primary tumor tissue

De-identified human prostate tissues were obtained from the UCSF BIOS tissue bank. Human prostate slices were prepared and cultured as previously published (37). Briefly, 8 mm diameter cores of putative benign and cancerous regions were taken from the peripheral zones according to gross analysis. The cores were aseptically cut to ∼300 mm thickness in the Krumdieck Tissue Slicer (Alabama Research and Development, Mundford, AL, USA). Five tissue slices were transferred to the titanium mesh inserts in 6-well plates containing 2.5 mL of complete PFMR-4A medium. Fully supplemented PFMR-4A media was kindly provided by Dr. Peehl at the UCSF Department of Radiology. The plates were incubated at 37°C in a humidified atmosphere containing 5% CO_2_ on a 30° angled rotating platform. After 48 hours, media was removed and kept on ice for exosome preparation.

### Exosome isolation and purification

To isolate exosomes from tumor cells, the cells were plated at a density of 3×10^6^ cells per 15 cm plate for MC38, TRAMP-C2, CT26, and B16F10 cells or 5×10^6^ cells per plate for PC3 and SK-MEL-28 cell lines and cultured in their complete media for 48 hours. To isolate exosomes from TRAMP-C2 cells for mass spectrometry, the cells were cultured in the complete media ± 10 ng/mL IFN-γ (Abcam, cat. Ab9922) for 48 hours prior to exosome collection. Culturing of primary tumor tissue slices was performed as described above. After culturing, exosomes were isolated from the media through differential ultracentrifugation by following our previously published protocol (5). In brief, pooled media from the cell or primary tissue culturing plates were spun at 300g for 10 minutes at room temperature to pellet cells. The supernatant was subsequently spun at 4°C at 2,000g for 20 minutes to pellet cell debris, 12,000g for 40 minutes to pellet microvesicles, and 100,000g for 70 minutes to pellet exosomes. The 100,000g pellet was resuspended in PBS and spun again at 100,000g for 70 minutes to wash the exosomes.

To purify exosomes from tumor cell lines or primary tissues for mass spectrometry, isolated exosomes were subjected to a sucrose density gradient. Exosomes were resuspended in 60% sucrose and loaded onto a gradient of 0%, 20%, 40% sucrose at increasing density. The gradient was spun at 4°C at 47,000 rpm for 16 hours. Sucrose fractions containing exosomes (20-40%) were identified via a refractometer, diluted with 1 mL PBS, and spun at 4°C at 100,000g (50,000 rpm) for 3 hours to pellet purified exosomes.

### Mass spectrometry

Isolated cell and exosome pellets from PC3, and SK-MEL-28 cells, and exosomes from primary tumors were processed and analyzed as previously described (38). Briefly, pellets were resuspended in chaotropic lysis buffer (50 mM Tris pH 8.5, 6M guanidinium hydrocholoride, 5 mM TCEP, and 10 mM chloroacetamide) and simultaneously lysed, reduced, and alkylated by heating at 97°C for 10 minutes with intermittent vortexing. Cell pellets were further disrupted using sonication. Samples were allowed to cool and insoluble debris removed by centrifugation (21,000g, 10 minutes). The resulting supernatant was diluted to 2 M guanidinium hydrocholoride with 50 mM Tris, pH 8.5 and protein concentration assessed by absorbance at 280 nm. Sequencing-grade trypsin (Promega) was added at a 1:100 ratio relative to total protein in the lysate and digestion allowed to proceed overnight at room temperature. After digest, samples were desalted using SOLA HRP SPE columns (ThermoFisher) following standard protocols. Eluted peptides were dried and resuspended in 0.1% formic acid with 2% acetonitrile. LC-MS/MS analysis was then performed on 1 μg of resuspended peptides using a Q Exactive Plus mass spectrometer as previously described (38, 39). Exosome samples from TRAMP-C2 cells were processed using the commercial Preomics iST kit and analyzed using a Bruker timsTOF Pro mass spectrometer as previously described (40). All proteomics data were evaluated using MaxQuant software. For data generated using the Q Exactive Plus mass spectrometer, data were analyzed, and label-free quantitation performed with MaxQuant version 1.5.1.2. Data were searched against the human proteome (SwissProt) with cysteine carbamidomethyl set as a fixed modification. N-terminal acetylation and methionine oxidation were set as variable modifications. Search results were filtered to a false discovery of 1% at both the peptide and protein levels. Data were visualized, processed, and compared using Perseus. Proteins with only one unique peptide were removed from analysis. LFQ intensities were log_2_(x) transformed and missing values imputed using standard Perseus settings (width of 0.3, downshift of 1.8). Statistical differences between cell and exosome protein content for PC3 and SK-MEL-28 lines were determined using student’s t-test. For analysis of the TRAMP-C2 samples, MaxQuant version 1.6.6.0 was used to search data against the mouse proteome (SwissProt) and similarly triaged using Perseus. For functional gene enrichment analysis, all proteins detected in at least one PC3 or SK-MEL-28 exosome samples were analyzed by ShinyGO 0.77 (http://bioinformatics.sdstate.edu/go/) to identify the immune regulation related molecules. String network of the functional and physical protein-protein interactions of the identified exosomal immune regulators was generated from Sting-db, Version 11.5 (https://string-db.org/) with a confidence score 0.4 and above (41).

### CRISPR/Cas9-mediated gene knockout

The sgRNA oligonucleotides (IDT DNA) targeting mouse *Pvrl2* or *Pvr* were cloned into pSpCas9(BB)-2A-GFP plasmid (PX458, ADDGENE). 6 μg of each vector was transfected into tumor cells plated on a 6-well plate using FuGENE6 transfection reagent (Promega, cat. E2691) and OPTI-MEM (GIBCO, ref. 31985-062). 48 hours post transfection, *Pvrl2* and *Pvr* KO clones were flow-sorted by GFP+ single cell cloning. After expansion, knockout clones were identified by flow cytometry analysis for cell surface expression of PVRL2 or PVR. MC38 *Pvrl2*; *Pvr* double KO cells were generated by transfecting the *Pvr* sgRNA containing PX458 plasmid into MC38 *Pvrl2* KO cells using the same strategy. MC38 *Pdl1* KO cells were generated through CRISPR Cas9-gRNA RNP-directed deletion by using a Lonza 4D-Nucleofector and a SF Cell Line 4D-Nucleofector X Kit S (Lonza, cat. V4XC-2032). In brief, 65 pmol of sgRNA targeting mouse *Pdl1* (IDT DNA) and 30 pmol of S.p. Cas9 Nuclease (IDT, cat. 1081058) were mixed and incubated at 37°C for 10 minutes to form the Cas9-sgRNA RNP complex. MC38 cells were suspended in 20 uL SF buffer with supplement and then the RNP mixture was added to the cell suspension for nucleofection. Following nucleofection, cells were allowed to recover for 3 days, and then the PD-L1 negative population was purified by three rounds of flow-sorting.

Mouse *Pvrl2* guide: GTCGGTGACAATCTGGACGG

Mouse *Pvr* guide: GAAATTCTTGGCTGCCCAAC

Mouse *Pdl1* guide: GTTTACTATCACGGCTCCAA (5)

### Western blot

Cell and exosome samples were lysed in RIPA buffer (ThermoFisher, cat. 89900) supplemented with PhosSTOP (Sigma-Aldrich, cat. 4906837001) and Complete Mini proteasome inhibitors (Sigma-Aldrich, cat. 05892791001). Total protein concentration was measured using BCA protein assay kit (ThermoFisher, cat. 23225). 30 μg total protein was loaded for PC3, TRAMP-C2, and MC38 cells and exosomes, and 40 μg for CT26 and B16F10 cells and exosomes. The cell and exosome protein samples were subjected to immunoblotting by following the manufacturer recommended protocols for the antibodies:

Primary antibodies: anti-mouse/human PVRL2 (EPR6717) (Abcam, cat. 135246), anti-α-Tubulin (Sigma-Aldrich, cat. T6074), anti-Hrs (C-7) (Santa Cruz, cat. sc-271455), anti-mouse/human PVRIG [EPR26274-202] (Abcam, cat. ab307595).

Secondary antibodies: goat anti-rabbit IgG (H+L) secondary antibody (ThermoFisher, cat. 35568), goat anti-mouse IgG (H+L) secondary antibody (ThermoFisher, cat. SA5-35521).

### NK cell cytotoxicity assay

NK cells were enriched from splenocytes from WT or *Pvrig* KO C57BL/6 mice by using the MojoSort™ Mouse NK Cell Isolation Kit (Biolegend, cat. 480049). Enriched NK cells were cultured and stimulated *in vitro* in RPMI-1640 Medium (ATCC, cat. 30-2001) containing 10% Fetal Bovine Serum (Corning, ref. 35-010-CV), Penicillin/Streptomycin (Sigma, cat. P4333), 1 mM sodium pyruvate (GIBCO, ref. 11360-070), 0.05 mg/mL Gentamicin (GIBCO, ref. 15750-060), 2-mercaptoethanol (GIBCO, car 21985-023), supplemented with 1,000 U/mL mouse IL-2 (PeproTech, cat. 212-12) for 7-9 days. Target cells (TRAMP-C2 WT and *Pvrl2* KO) were labeled with CellTrace Violet (ThermoFisher, cat. C34557), and co-cultured with WT or *Pvrig* KO NK cells as effectors at 1:1 ratio for 4 hours. Then the NK cell lysis was analyzed by flow cytometry with propidium iodide staining (Invitrogen, cat. P3566) according to the published protocol (42).

To generate *Pvrig*; *Tigit* and *Pvrig*; *Cd96* double KO NK cells, *Pvrig* KO NK cells were isolated from spleens of *Pvrig* KO mice and transfected with two sgRNAs targeting mouse *Tigit* or *Cd96* (IDT DNA) using Cas9-gRNA RNP-directed gene deletion on day 5. The nucleofection was conducted using a Lonza 4D-Nucleofector and a P3 Primary Cell 4D-Nucleofector X Kit S (Lonza, cat. V4XP-3032) with the same protocol described earlier. Subsequently, TIGIT and CD96 negative cell populations were purified by flow-sorting on day 8. Cytotoxicity assays were performed on day 11 using the same protocol above.

Mouse *Tigit* guides: CTGAAGTGACCCAAGTCGAC; TTCAGTCTTCAGTGATCGGG

Mouse *Cd96* guides: GATGACGTGTATGCTCTACC; TCCAAATCCAAGACGATGGC

### Tumor cell injections

Moues tumor cells were cultured in their regular growth media as mentioned above. Prior to injection, the cells were harvested by trypsinization, washed once with PBS, and then resuspended in PBS (1×10^6^ cells/100 μL). 1×10^6^ cells per mouse were injected subcutaneously in the right flank of age matched (8-11-week-old) male mice. Mice were considered “end stage” when the tumor volume reached 1,500 mm^3^ or the tumor became ulcerated. Tumor growth was monitored every 2 or 3 days by measuring tumor length and width using caliper. Tumor volume was calculated according to the equation: 0.5 × (width)^2^ × length.

### Mouse treatments

Exosome injection: Exosomes were isolated from cultured WT and *Pvrl2* KO MC38 cells as described above. 100kg pellet was resuspended in PBS (100 μL/15 cm plate), and 100 μL was injected into the tail vein of age matched (8-11-week-old) male WT C57BL/6 mice. On the same day, 1×10^6^ *Pvrl2* KO MC38 cells in 100 μL of PBS were injected subcutaneously in the right flank of the mice to establish tumors. Exosome isolation and injection was repeated three times a week for two weeks.

Antibody injection: For immune cell depletion, anti-mouse CD8α (2.43) (Bio X Cell, cat. BE0061) and rat IgG2b isotype control (LTF-2) (Bio X Cell, cat. BE0090) antibodies were diluted to 100 μg/100 μL with pH 7.0 dilution buffer (Bio X Cell, cat. IP0070); anti-mouse CD4 (GK1.5) (Bio X Cell, cat. BE0003-1) was diluted to 200 μg/100 μL with pH 6.5 dilution buffer (Bio X Cell, cat. IP0065); anti-mouse NK1.1 (PK136) (Bio X Cell, cat. BE0036) was diluted to 200 μg/100 μL with pH 7.0 dilution buffer (Bio X Cell, cat. IP0070). Then the antibodies were intraperitonially (I.P.) injected into mice (100 μL/mouse) starting one day prior to tumor injection, followed by two more weekly doses. For TIGIT blockade, anti-mouse TIGIT (1B4) (Absolute Antibody, cat. Ab01258-1.1) (1 mg/mL) or mouse IgG1 isotype control (MOPC-21) (Bio X Cell, cat. BE0083) were intraperitonially (I.P.) injected into mice starting on day 4 post tumor injection followed by a serial 4 doses (200 μg/mouse) every three days, and a maintenance dose (100 μg/mouse) on day 20.

### Immune-profiling

Age match and randomly assigned male C57BL/6 mice were implanted subcutaneously (S.C.) with 1×10^6^ MC38 WT or *Pvrl2* KO cells to the right flank. On day 25, mice were euthanized, and tumors surgically removed with sterilized surgical equipment, weighed, and minced into small pieces using scissors. The minced tumor tissue was transferred to a 6-well plate containing 3 mL/well of tumor digestion media (NK cell media + 1 mg/mL collagenase IV (Sigma, cat. C5138) + 0.2 mg/mL DNAse I (Roche, cat. 10104159001) and incubated on a shaker at 37°C for 1 hour. Cell mixtures were then filtered through a 70 μm strainer into 50 mL conical tubes. Cells were then washed once with FACS buffer (PBS + 2% heat inactivated FBS + 1 mM EDTA) and counted. CD45+ cells were enriched using the EasySep Mouse TIL (CD45) positive selection kit (STEMCELL, cat. 100-0350) by following the manufacturer’s protocol.

Single cell suspensions (1×10^6^ cells) were first stained with cell viability dye (1:1000 in PBS) (eBioscience, cat. 65-0866-14; CYTEK, cat. SKU 13-0865-T100) for 30 minutes at 4°C in dark to exclude dead cells. After two washes with FACS buffer, cells were incubated with Fc Block (Tombo Biosciences, cat. 70-0161) for 10 minutes and then co-incubated with fluorescently labelled antibodies for 30 minutes at 4°C in dark, followed by three washes with FACS buffer. Detailed information of the flow cytometry antibodies used in this study are listed below. Flow cytometry was performed on LSRII S854-S864, and data was analyzed by FlowJo 10.8.1.

Flow antibodies:

Anti-mouse CD8 Brilliant Violet 605 (53-6.7) (Biolegend, cat. 100744),

anti-mouse CD4 Brilliant Violet 421 (GK1.5) (Biolegend, cat. 100437),

anti-mouse NK1.1 PE (PK136) (Biolegend, cat. 108707),

anti-mouse NK1.1 Brilliant Violet 711 (PK136) (Biolegend, cat. 108745),

anti-mouse CD107a (LAMP-1) Brilliant Violet 711 (1D4B) (Biolegend, cat. 121631),

anti-mouse PD-1 (CD279) APC (J43) (BD Biosciences, cat. 562671),

anti-mouse PVRL2 Brilliant Violet 421 (BD Biosciences, cat. 748046),

anti-mouse PVR PE (TX56) (Biolegend, cat. 131507),

anti-mouse PVR APC (TX56) (Biolegend, cat. 131509),

anti-mouse TIGIT PE (1G9) (Biolegend, cat. 142104),

anti-mouse CD96 APC (3.3) (Biolegend, cat. 131711),

anti-mouse DNAM-1 Brilliant Violet 785 (TX42.1) (Biolegend, cat. 133611),

anti-mouse CD98 PE (4F2) (Biolegend, cat. 128207),

anti-mouse PD-L1 Super Bright 780 (MIH5) (Invitrogen, cat. 78-5982-82),

Brilliant Violet 605 Rat IgG2a, k Isotype control (RTK2758) (Biolegend, cat. 400539),

Brilliant Violet 421 Rat IgG2b, κ Isotype control (RTK4530) (Biolegend, cat. 400639),

PE Mouse IgG2a, κ Isotype control (MOPC-173) (Biolegend, cat. 400211),

Brilliant Violet 711 Rat IgG2b, κ Isotype control (RTK4530) (Biolegend, cat. 400653),

APC Hamster IgG2, κ Isotype control (B81-3) (BD Biosciences, cat. 562169).

### Immunofluorescence and image analysis

Age matched and randomly assigned male C57BL/6 mice were implanted subcutaneously (S.C.) with 1×10^6^ MC38 WT or *Pvrl2* KO cells. On day 25 or 27, mice were euthanized, and tumors dissected and rinsed with PBS. Tumors were fixed in 10% neutral buffered formalin at room temperature overnight, after which they were dehydrated by sucrose (5% for 1 hour, 10% for 1 hour, 20% overnight) and embedded in the Scigen Tissue-Plus™ O.C.T. Compound (ThermoFisher, cat. 23-730-571).

For immunofluorescence, tumors were sectioned into 10 μm slides using a Leica CM 3050S cryostat. Sections were rehydrated with PBS for 10 minutes. Antigen retrieval was performed using EDTA antigen retrieval buffer (10 mM Tris base, 1 mM EDTA solution, 0.05% Tween 20, pH∼9) in a steamer for 20 minutes. Sections were incubated in blocking buffer (PBS with 1% BSA and 5% donkey serum) at room temperature for 1 hour before overnight incubation with primary antibodies diluted 1:100 in blocking buffer at 4°C. Anti-CD8a (Invitrogen, cat. 14-0808-80) (1:100) and anti-NCR1 (Abcam, cat. ab233558) (1:100) primary antibodies were used. After primary body incubation, sections were washed with TBST (0.05% Tween-20 in TBS) and then incubated with DAPI (Invitrogen, cat. D1306) (1:1000) and fluorescence-conjugated secondary antibodies diluted 1:200 in blocking buffer for 1 hour at room temperature. Donkey anti-Rat IgG (H+L) Alexa Fluor™ 488 (Invitrogen, cat. A21208) and Goat anti-Rabbit IgG (H+L) Alexa Fluor™ 594 (Invitrogen, cat. A11012) were used for CD8a and NCR1 staining, respectively. Sections were washed with TBST and mounted using the ProLong™ Gold Antifade Mountant (Invitrogen, cat. P36930). Images were acquired using a Leica SP8 confocal microscope. All images were processed with ImageJ 1.53.

A 3-6 μm z-stack with system optimized step-size was taken for each field of view. To quantify the total number of CD8 T cells and NK cells in each z-stack of field of view, maximum intensity projection was applied to all slices in the z-stack. CD8 T cells and NK cells were then counted manually. A representative slice in the z-stack was shown.

### Statistical analysis

All statistical analyses were processed with GraphPad Prism, Version 9.4.1 (GraphPad). Statistical significance between groups of areas under curves (AUC) of tumor growth, *in vitro* NK cytotoxicity assay, immune-profiling, and IF images were calculated using unpaired student’s t test. Statistical significance for mouse survival was analyzed by log rank test. No statistical method was used to predetermine sample size. No data points were excluded from the analyses of all experiments. In all cases, significance was defined by a *P* value of 0.05 and below. Details regarding the *P* values, number of replicates and the definition of center and error bars are indicated in figures and figure legends. *P* values for AUC comparisons not shown in the figures can be found in Supplementary Table S1.

### Data and material availability

The mass spectrometry proteomics data that support the findings of this study have been deposited to the ProteomeXchange Consortium via the PRIDE (43) partner repository with the dataset identifier PXD044245. All the remaining data that support the conclusions from this study are included in this article. *Pvrl2* and *Pvr* KO tumor cell lines and *Pvrig* KO mice generated in this study will be made available on request with completed Material Transfer Agreements. Further information and data are available from the corresponding author on reasonable request.

## Results

### Proteomic analysis identifies PVRL2 on tumor-derived exosomes

To identify immunosuppressive molecules beyond PD-L1 that are present on tumor-derived exosomes, we employed mass spectrometry-based proteomic analysis on exosomes isolated from two human cancer cell lines, PC3 (prostate cancer) and SK-MEL-28 (melanoma) (Fig. 1A). The analysis revealed over two thousand proteins on PC3 and SK-MEL-28 exosomes, including 78 proteins identified as regulators of the immune responses, as determined by functional gene enrichment analysis (Supplementary Fig. S1A). We extended our proteomic analysis to the mouse prostate cancer cell line TRAMP-C2 and two *ex-vivo* cultured primary human prostate cancer tumor slices (Fig. 1A). Among the list of 78 immune regulators from PC3 and SK-MEL-28 exosomes, 28 proteins were also detected in at least one sample derived from TRAMP-C2 and primary tumor exosomes (Fig. 1B; Supplementary Fig. S1B). We evaluated for proteins enriched on exosomes relative to their cell of origin uncovering 274 proteins from PC3 and 405 proteins from SK-MEL-28 exosomes including 4 out of the 28 shared immunoregulators that were significantly enriched in the PC3 and SK-MEL-28 exosomes relative to the cells (> 2-fold, *P* < 0.05) (Fig. 1C and D; Supplementary Fig. S1C). These four proteins were DLG1, YES1, PVRL2 and CTNNB1 (Fig. 1C and D; Supplementary Fig. S1C). We focused on PVRL2 given its previously reported immunoregulatory functions (14, 18, 25, 44). To validate and expand on our proteomic data, we performed immunoblot analysis for PVRL2 on cell and exosome fractions from TRAMP-C2 and PC3 as well as three other mouse syngeneic models including two colorectal cell lines (MC38 and CT26) and a melanoma cell line (B16F10). All five lines showed robust levels of PVRL2 protein in both cells and exosomes (Fig. 1E; Supplementary Fig. S1D). These data uncover the presence of PVRL2 in both the cellular and exosome fractions of multiple mouse and human tumor models representing different cancer types.

**Figure 1:**
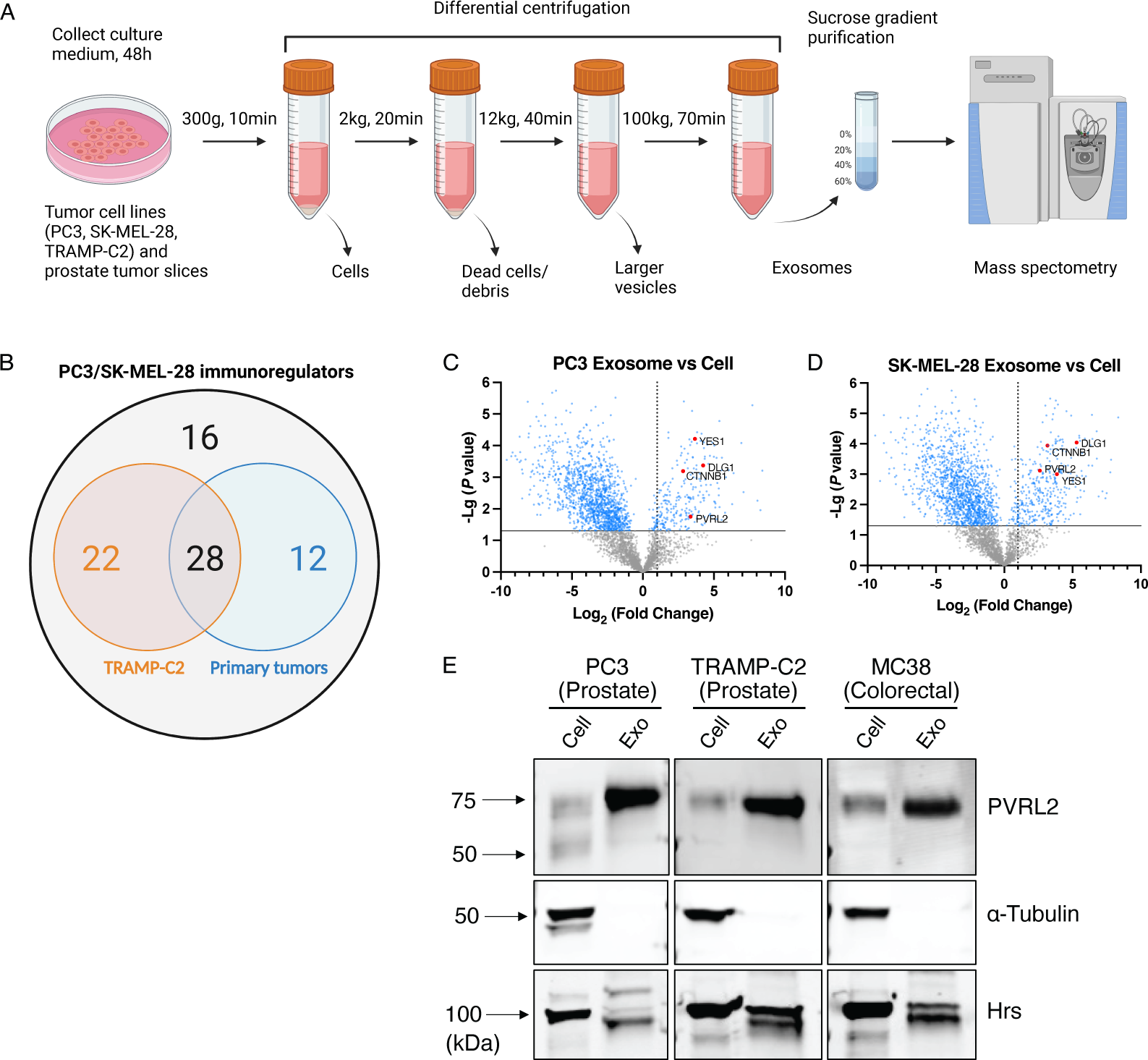
Proteomic analysis identifies PVRL2 on tumor-derived exosomes. **(A)** Schematic of exosome collection: exosomes were collected from the indicated tumor cell lines and primary tumor slices via differential centrifugations and purified by sucrose gradient. Exosomes and cells were then lysed, and their respective proteomes were analyzed by mass spectrometry. **(B)** The numbers of shared immunoregulatory molecules identified from mass spectrometry results of the exosomes from the indicated tumor cell lines and primary tumors. **(C-D)** Volcano plots present protein abundance differences in exosomes over in cells in PC3 (C) and SK-MEL-28 (D) cell lines as determined by label free quantitation. Proteins on the right of the volcano plot represent proteins enriched in exosomes versus cells. Proteins with a log_2_ (fold change) value > 1 are over 2-fold enriched. **(E)** Western blot analysis for PVRL2 in the cells and exosomes (exo) from the indicated tumor cell lines. 30 μg of total protein was loaded for each sample. A-tubulin was used as the loading control for cells, and Hrs as the loading control for exosomes.

### PVRL2 promotes tumor growth through an immune-dependent mechanism

Given the presence of PVRL2 on both tumor exosomes and cells, we performed functional experiments in mice to test the relevance of PVRL2 in the regulation of anti-tumor immune response. We used CRISPR/Cas9-directed mutagenesis to knockout *Pvrl2* (gene encoding PVRL2, also known as *Nectin2*) in the four mouse syngeneic tumor cell lines (Supplementary Fig. S2A). Flow cytometry analysis confirmed loss of the PVRL2 protein in the mutant clones of each line (Supplementary Fig. S2B-S2E). *In vitro* growth analysis showed no effect of PVRL2 loss on tumor cell growth rate *in vitro* (Supplementary Fig. S2F-S2I). The mutant cell lines were transplanted into immunocompetent isogenic mice (C57BL/6 for MC38, TRAMP-C2, and B16F10; BALB/cJ for CT26). In all four models, PVRL2 loss led to a dramatic reduction in tumor growth and extended the survival of the mice (Fig. 2A-H; Supplementary Fig. S2J-S2M). Comparison to a *Pdl1* knockout in the MC38 model showed an even greater impact on tumor growth with PVRL2 loss than with PD-L1 loss (Supplementary Fig. S2N-S2Q). To determine whether the PVRL2 effects were specifically through the regulation of the immune response, we repeated the experiments in NOD CRISPR *Prdkc Il2r Gamma* (NCG) mice, which are deficient for T, B, and NK cells and have reduced macrophage and dendritic cell function (45–47). In this background, the wild-type (WT) and *Pvrl2* knockout (KO) tumors grew equally fast resulting in the rapid demise of their hosts (Fig. 2I-P). Together, these results identify PVRL2 as a key promoter of tumor growth *in vivo*, functioning through an immune-dependent mechanism.

**Figure 2:**
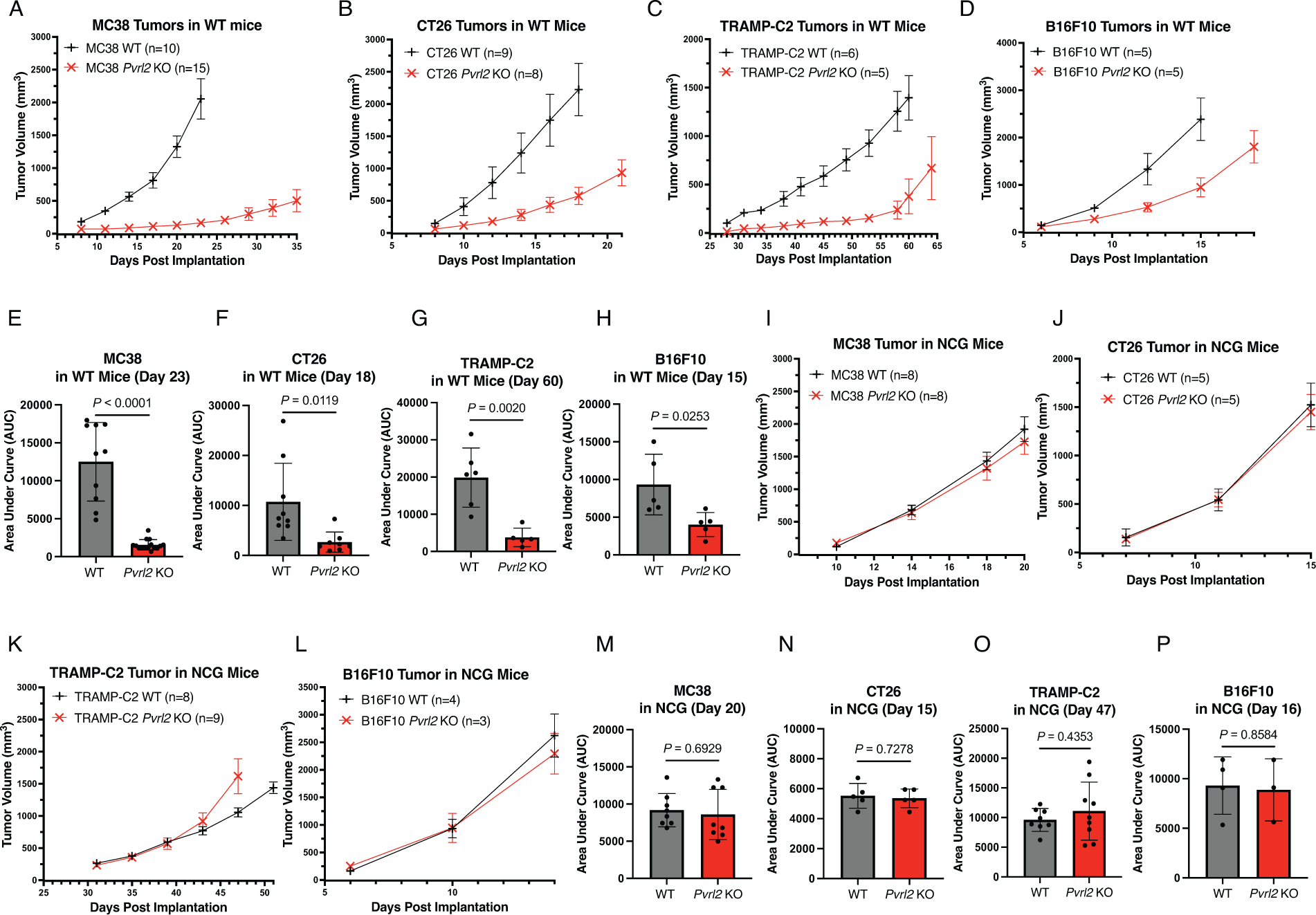
PVRL2 promotes tumor growth through an immune-dependent mechanism. **(A-D)** Average tumor volume over time following subcutaneous injection of 1 × 10^6^ wild-type (WT) and *Pvrl2* knockout (KO) MC38 (A), TRAMP-C2 (C), and B16F10 (D) cells in C57BL/6 mice, and CT26 (B) in BALB/cJ mice. Error bars represent standard error of the mean (SEM). **(E-H)** Area under the curves (AUC) of the MC38 (E), CT26 (F), TRAMP-C2 (G), and B16F10 (H) tumors from (A-D) calculated at day when the first mouse reached end point: day 23, 18, 60, and 15, respectively. Dots represent individual mice. *P* values are calculated by unpaired t test. Error bars represent standard deviation (SD). **(I-L)** Average tumor volume over time following subcutaneous injection of 1 × 10^6^ MC38 (I), CT26 (J), TRAMP-C2 (K), and B16F10 (L) WT and *Pvrl2* KO cells in NCG mice. Error bars represent SEM. **(M-P)** Area under the curves of the MC38 (M), CT26 (N), TRAMP-C2 (O), and B16F10 (P) tumors from (I-L) calculated at day when the first mouse reached end point: day 20, 15, 47, and 16, respectively. Dots represent individual mice. *P* values are calculated by unpaired t test. Error bars represent SD.

### Exosomal PVRL2 partially rescues the phenotype of *Pvrl2* KO tumors

We addressed whether and to what degree PVRL2 secreted in exosomes contributes to the overall ability of PVRL2 to promote tumor growth. To address this question, we focused on the rapidly growing MC38 model. *Pvrl2* KO MC38 cells were transplanted into WT C57BL/6 mice. Exosomes were isolated from cultured WT and *Pvrl2* KO MC38 cells. These exosomes were injected into the tail vein of the mice starting on the same day as tumor cell transplant and continued three times per week for two weeks (Fig. 3A). The injection of exosomes from the WT cells significantly accelerated the growth of *Pvrl2* KO tumors and reduced the survival of the treated mice (Fig. 3B-D). However, growth remained well below that of transplanted WT cells (Fig. 3E). In contrast, *Pvrl2* KO exosomes did not significantly impact tumor growth or mouse survival (Fig. 3B-D). These findings show that exosomal PVRL2 can act to partially rescue growth of *Pvrl2* KO tumor cells. However, exosomal PVRL2’s contribution to promoting tumor growth appears small relative to that of cellular PVRL2.

**Figure 3:**
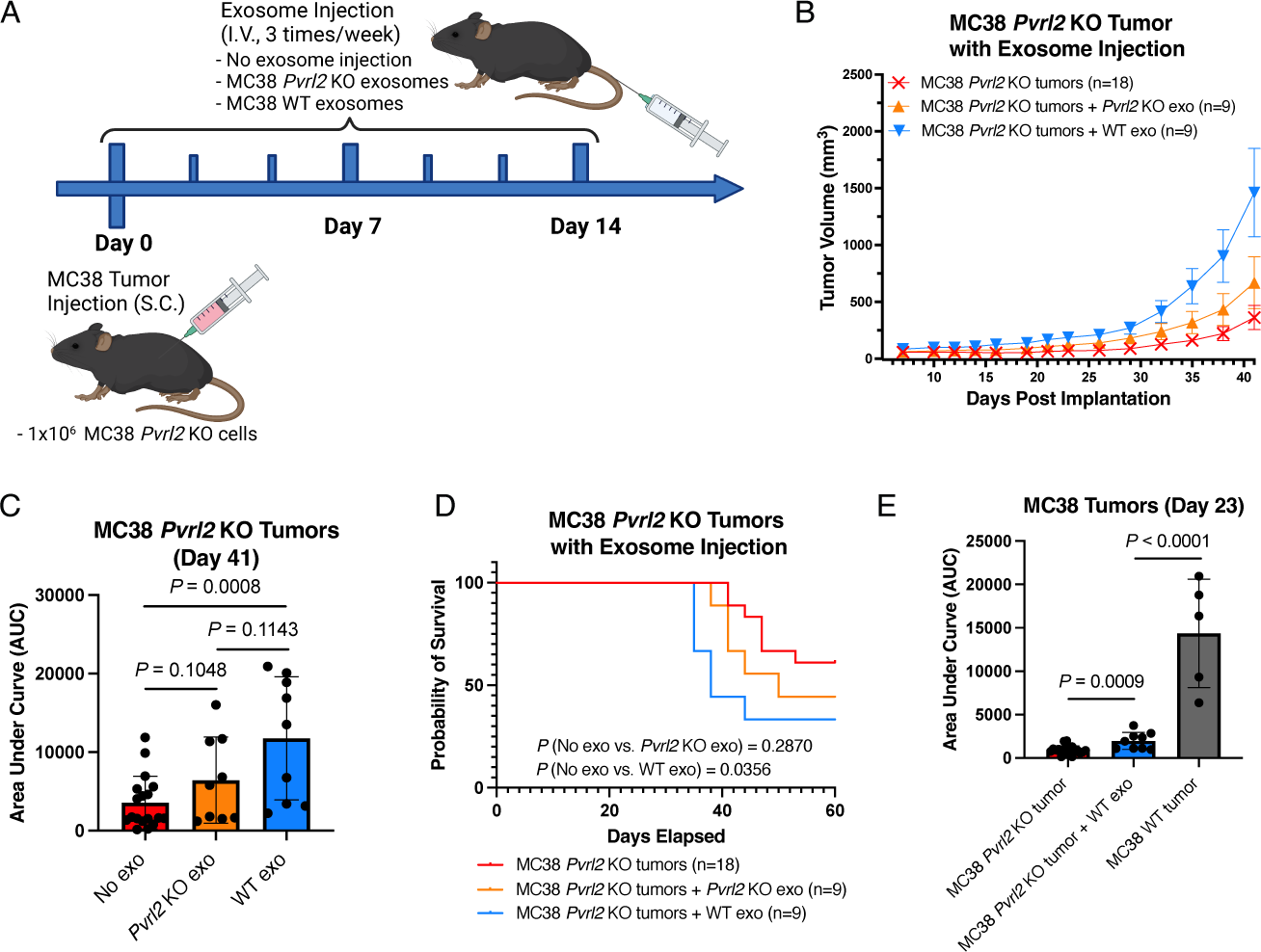
Exosomal PVRL2 partially rescues the phenotype of *Pvrl2* KO tumors. **(A)** Schematic of the experiment: 1 × 10^6^ MC38 *Pvrl2* KO cells were injected into C57BL/6 mice, and starting from the same day, exosomes collected from MC38 WT and *Pvrl2* KO cells were injected into the mice through tail vein according to the indicated timeline. **(B)** Average tumor volume over time following the injection of MC38 *Pvrl2* KO tumors along with no exosome injection, MC38 WT exosomes, and MC38 *Pvrl2* KO exosomes as indicated in (A). Error bars represent SEM. **(C)** Area under the curves of the tumor growth in (B) on day 41. Dots represent individual mice. *P* values are calculated by unpaired t test. Error bars represent SD. **(D)** Mouse survival curves following injections as described in (B). *P* values are calculated by log rank test. **(E)** Area under the curves of the growth of MC38 *Pvrl2* KO tumors with no exosome injection or WT exosome injection from (B), in comparison with the MC38 WT tumors from Fig. 2 (A) on day 23. Dots represent individual mice. *P* values are calculated by unpaired t test. Error bars represent SD.

### PVRL2 regulates CD8 T cell and NK cell activation

The PVRL2 receptor PVRIG is expressed on T cells and NK cells (48), and has been shown to display immune inhibitory function on these cells in both mouse and human models (19, 24, 25, 49-51). However, there is currently no direct evidence showing that PVRL2 regulates these specific cell populations. Therefore, we set out to resolve what immune populations were impacted and responsible for PVRL2’s role in promoting tumor growth. First, to determine whether PVRL2 primarily functions through adaptive or innate immunity, we transplanted MC38 tumor cells into *Rag1* KO mice, which lack functional T and B cells and thus the adaptive immune response. Interestingly, even in the absence of adaptive immunity, MC38 WT tumors maintained significantly faster growth than *Pvrl2* KO tumors, indicating that the remaining innate immune response plays a crucial role in PVRL2 function (Fig. 4A and B). However, the ratios of WT over KO tumor size and the areas under the curve (AUC) were reduced in the *Rag1* KO mice supporting a role of the adaptive response as well (Supplementary Fig. S3A).

**Figure 4:**
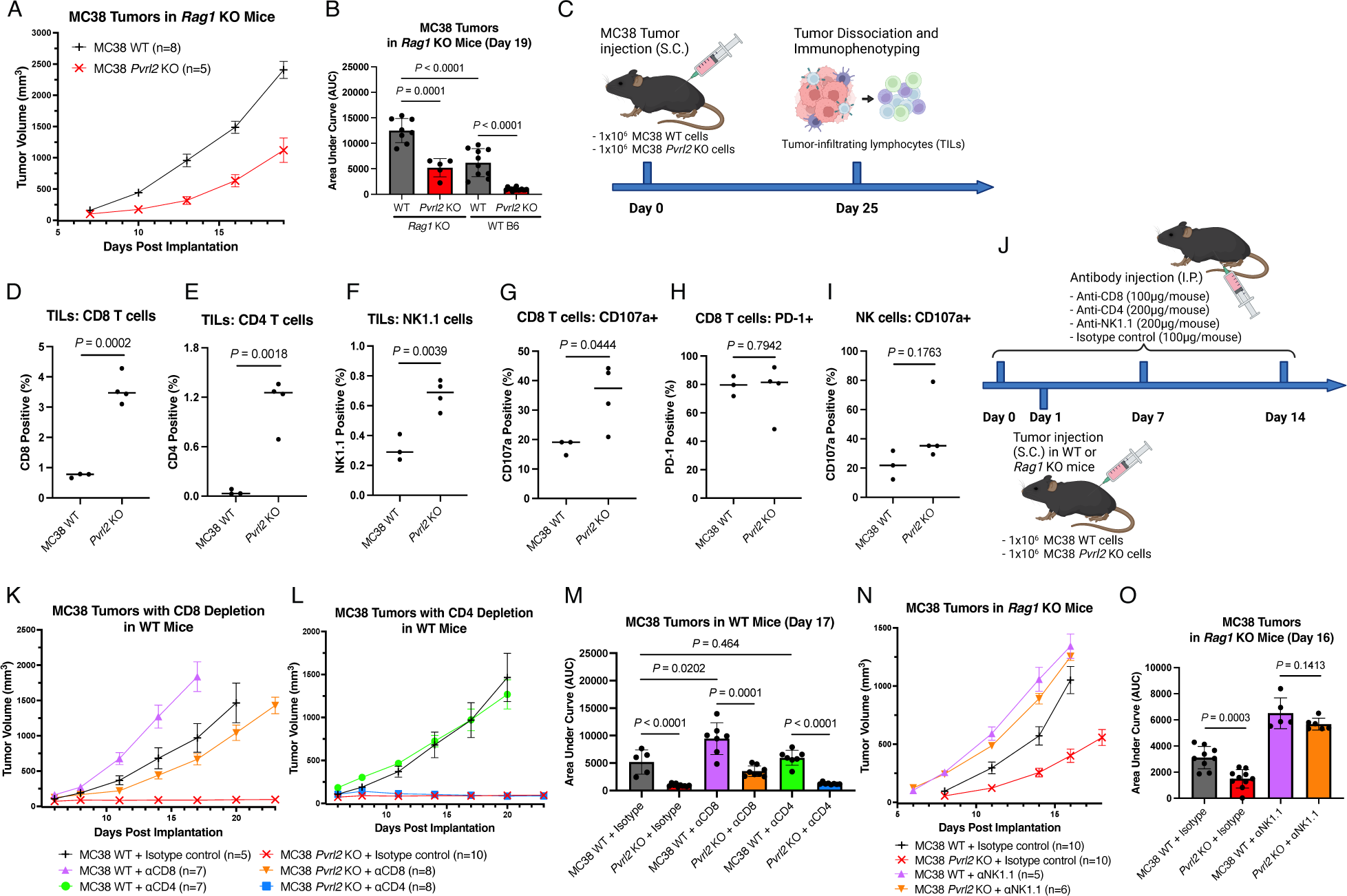
PVRL2 regulates CD8 T cell and NK cell activation. **(A)** Average tumor volume over time following subcutaneous injection of 1 × 10^6^ WT and *Pvrl2* KO MC38 cells in *Rag1* KO mice. Error bars represent SEM. **(B)** Area under the curves of the MC38 tumors from (A) and Fig. 2 (A) on day 19. Dots represent individual mice. *P* values are calculated by unpaired t test. Error bars represent SD. **(C)** Schematic of immunophenotyping experiment: 25 days after subcutaneous injection of 1 × 10^6^ MC38 WT and *Pvrl2* KO cells in C57BL/6 mice, the tumors were collected. CD45+ cell populations were isolated from the tumor dissociates using CD45 positive magnetic selection kit. Then the isolated cells were subjected to viability dye, CD8, CD4, NK1,1, CD107a, and PD-1 staining followed by flow cytometry analysis. **(D-F)** Flow cytometric quantification of the percentage of CD8+ (D), CD4+ (E), and NK1.1+CD8-(NK cells) (F) populations, respectively, in the CD45+ cells isolated from the MC38 WT (n = 3) and *Pvrl2* KO (n = 4) tumors as indicated in (C). *P* values are calculated by unpaired t test. Line represents mean. **(G-H)** Quantification of the percentage of CD107a+ (G) and PD-1+ (H) cells among the CD8 T cell (D) population. *P* values are calculated by unpaired t test. Line represents mean. **(I)** Quantification of the percentage of CD107a+ cells among the NK cell (F) population. *P* value is calculated by unpaired t test. Line represents mean. **(J)** Schematic of experiment design in (K-O): After subcutaneous injection of 1 × 10^6^ MC38 WT and *Pvrl2* KO cells in C57BL/6 WT or *Rag1* KO mice, the mice were treated with anti-CD8, CD4 or NK1.1 depleting antibodies or isotype control at the indicated serial doses and schedule. **(K-L)** Average tumor volume over time following subcutaneous injection of 1 × 10^6^ MC38 WT and *Pvrl2* KO cells in WT C57BL/6 mice with anti-CD8 (K) and anti-CD4 (L) depleting antibodies or isotype control as indicated in (J). Error bars represent SEM. **(M)** Area under the curves of the MC38 tumors from (K) and (L) on day 17. Dots represent individual mice. *P* values are calculated by unpaired t test. Error bars represent SD. **(N)** Average tumor volume over time following subcutaneous injection of 1 × 10^6^ MC38 WT and *Pvrl2* KO cells in *Rag1* KO mice with anti-NK1.1 depleting antibody or isotype control as indicated in (J). Error bars represent SEM. **(O)** Area under the curves of the MC38 tumors from (N) on day 16. Dots represent individual mice. *P* values are calculated by unpaired t test. Error bars represent SD.

To understand which specific immune populations were responsible for PVRL2’s regulation of adaptive and innate immunity, we performed immune profiling of the tumor microenvironment (TME) by flow cytometry (Fig. 4C). Isolated CD45+ cells from MC38 WT and *Pvrl2* KO tumors were stained with a viability dye and antibodies against CD8, CD4, NK1.1, and the activation markers CD107a and PD-1 (Supplementary Fig. S3B-S3D). *Pvrl2* KO tumors had significantly higher fractions of CD8, CD4 T cells, and NK cells (Fig. 4D-F). Furthermore, a significantly larger proportion of CD8 T cells expressed the degranulation marker CD107a in *Pvrl2* KO tumors, indicating enhanced activation of these cells (Fig. 4G; Supplementary Fig. S3C). In contrast, PD-1 was unchanged (Fig. 4H; Supplementary Fig. S3D). NK cells in *Pvrl2* KO tumors also exhibited slightly higher CD107a positivity, although the data did not reach statistical significance (Fig. 4I). These findings show that *Pvrl2* KO promotes the infiltration and activation of adaptive and innate immune cells. Immunofluorescence (IF) staining of the tumor sections confirmed the increased infiltration of CD8 T cells and NK cells in MC38 *Pvrl2* KO tumors compared to WT tumors (Supplementary Fig. S3E-S3H).

To further validate that PVRL2’s function is dependent on T cells and NK cells *in vivo*, we depleted these specific cell populations with antibodies to CD8, CD4 or NK1.1 (Fig. 4J). Depletion of CD8 T cells resulted in a significant promotion of both MC38 *Pvrl2* KO and WT tumor growth (Fig. 4K and M) and extended the survival of the mice (Supplementary Fig. S3J). Notably, although the WT tumors still exhibited significantly faster growth than *Pvrl2* KO tumors, the difference between WT and *Pvrl2* KO tumors became smaller upon CD8 T cell depletion, as indicated by the ratio of WT tumor over *Pvrl2* KO tumor size and the AUC (Supplementary Fig. S3I). In contrast, CD4 T cell depletion did not impact tumor growth or survival (Fig. 4L and M; Supplementary Fig. S3J). To specifically measure the contribution of NK cells, we depleted NK cells in the *Rag1* KO background. This led to the abolishment of any differences between MC38 *Pvrl2* KO and WT tumor growth and mouse survival (Fig. 4N and O; Supplementary Fig. S3K). Together, these results show that PVRL2 on tumor cells suppresses both adaptive and innate immune responses and does so through inhibition of CD8 T and NK cells, respectively.

### PVRL2 functions through a PVRIG independent mechanism

As PVRL2 has been shown to bind to the co-inhibitory receptor PVRIG, it is assumed that PVRIG underlies PVRL2 function in the suppression of antitumor immune response (19, 25, 49-51). However, no evidence exists indicating whether PVRL2 directly regulates PVRIG. Our data shows a much greater effect of PVRL2 loss on tumor growth than previously published for PVRIG loss or antibody blockade (24, 51). To ask whether PVRIG is indeed the primary receptor responsible for mediating PVRL2’s ability to promote tumor growth, we produced *Pvrig* KO mice using sperm from the Knockout Mouse Project (KOMP). The knockout involves deletion spanning exons 1 through 4 plus part of 5 removing the entire coding region of *Pvrig* in a C57BL/6 background (Supplementary Fig. S4A). Genotyping by PCR validated the loss of *Pvrig* (Supplementary Fig. S4A-S4C). Furthermore, western blot analysis confirmed the depletion of PVRIG protein in the splenocytes from *Pvrig* KO mice (Supplementary Fig. S4D). All three WT and *Pvrl2* KO C57BL/6 mouse syngeneic mouse models were then transplanted into the mice. Surprisingly, the removal of PVRIG did not influence the growth or survival of WT MC38 tumors (Fig. 5A-C). Furthermore, the removal of PVRL2 was equally effective in both inhibiting MC38 tumor growth and enhancing survival in WT and *Pvrig* KO backgrounds (Fig. 5A-C). This effect cannot be ascribed to the indirect regulation of other Nectin family immunoregulatory receptors, as the loss of PVRIG in mice did not lead to significant alterations in the expression of these receptors, including TIGIT, DNAM-1, and CD96, on CD8 T cells and NK cells (Supplementary Fig. S4E and S4F). Similar results were observed with the TRAMP-C2 model, where PVRIG loss had no impact on the growth of WT or *Pvrl2* KO tumors (Fig. 5D-F). However, in the B16F10 model, the loss of PVRIG resulted in a notable, albeit partial, decrease in tumor growth (Fig. 5G and H); survival showed a trend towards improvement, but was not statistically significant (Fig. 5I). Importantly, the loss of PVRL2 alone had a greater impact compared to PVRIG loss alone, and PVRL2 loss further reduced tumor growth in a *Pvrig* KO background (Fig. 5G and H). In contrast, the growth of *Pvrl2* KO tumors was not additionally affected in the *Pvrig* KO background across all three cell lines (Fig. 5A-I). These data show that PVRL2 functions beyond PVRIG in promoting tumor growth.

**Figure 5:**
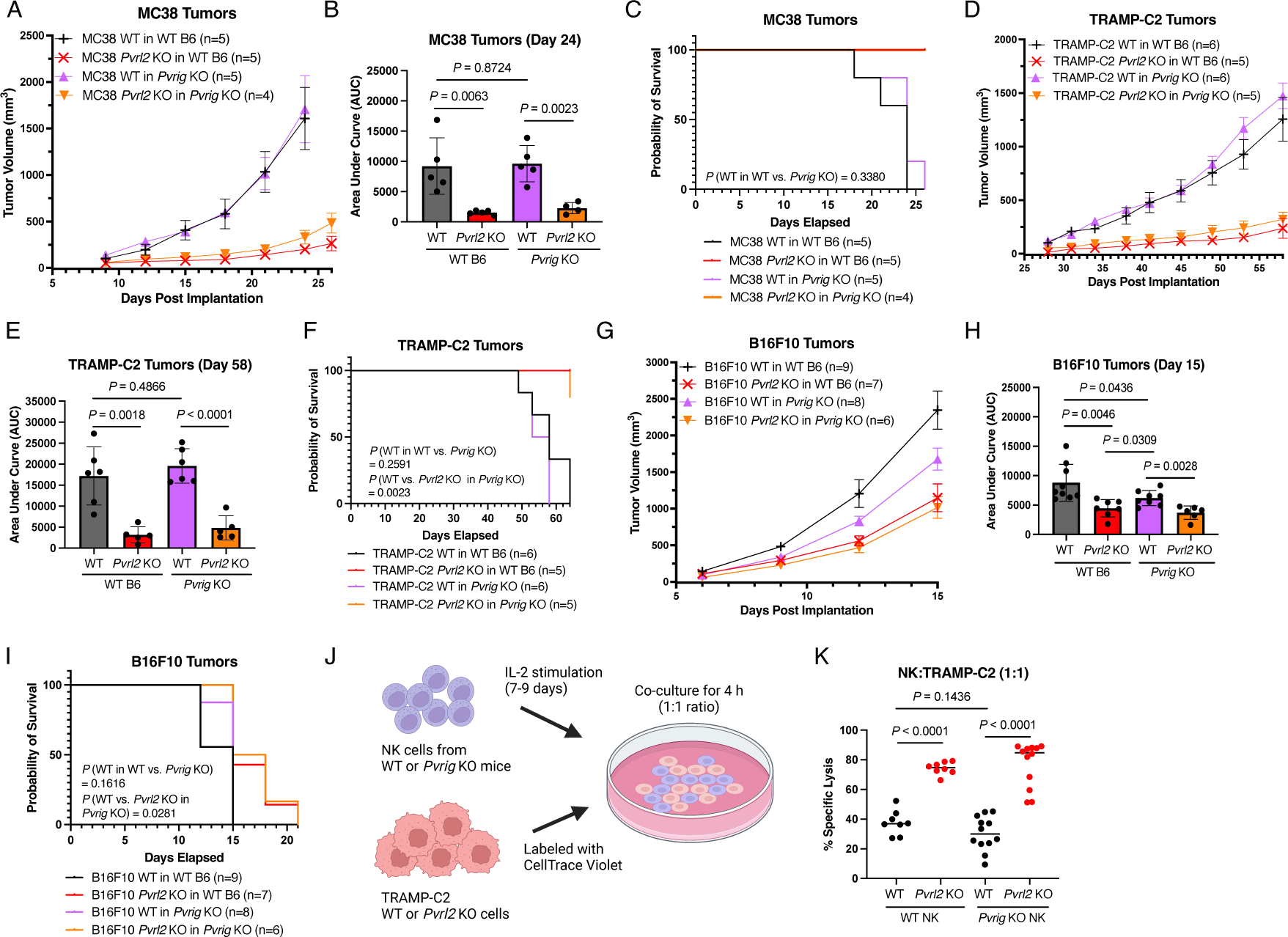
PVRL2 functions through a PVRIG independent mechanism. **(A)** Average tumor volume over time following subcutaneous injection of 1 × 10^6^ MC38 WT and *Pvrl2* KO cells in WT C57BL/6 mice and *Pvrig* KO mice. Error bars represent SEM. **(B)** Area under the curves of the MC38 tumors from (A) on day 24. Dots represent individual mice. *P* values are calculated by unpaired t test. Error bars represent SD. **(C)** Mouse survival curves following subcutaneous injections as described in (A). *P* values are calculated by log rank test. **(D)** Average tumor volume over time following subcutaneous injection of 1 × 10^6^ TRAMP-C2 WT and *Pvrl2* KO cells in WT C57BL/6 and *Pvrig* KO mice. Error bars represent SEM. **(E)** Area under the curves of the TRAMP-C2 tumors from (D) on day 58. Dots represent individual mice. *P* values are calculated by unpaired t test. Error bars represent SD. **(F)** Mouse survival curves following injections as described in (D). *P* values are calculated by log rank test. **(G)** Average tumor volume over time following subcutaneous injection of 1 × 10^6^ B16F10 WT and *Pvrl2* KO cells in WT C57BL/6 mice and *Pvrig* KO mice. Error bars represent SEM. **(H)** Area under the curves of the B16F10 tumors from (G) on day 15. Dots represent individual mice. *P* values are calculated by unpaired t test. Error bars represent SD. **(I)** Mouse survival curves following subcutaneous injections as described in (G). *P* values are calculated by log rank test. **(J)** Schematic of experiment design in (K): NK cells were isolated from WT C57BL/6 or *Pvrig* KO mice. After 7-9 days *in vitro* stimulation with IL-2, NK cells were co-cultured at 1:1 TRAMP WT or *Pvrl2* KO tumor cells for 4 hours. Then NK cell lysis was evaluated by live/dead cell dye followed by flow cytometry. **(K)** Percentage lysis of TRAMP-C2 WT and *Pvrl2* KO cells after co-culturing with WT or *Pvrig* KO NK cells at 1:1 ratio. Dots represent individual replicates. 4 replicates per condition in each experiment (n=2 and n=3 independent experiments for WT NK cells and *Pvrig* KO NK cells, respectively). *P* values are calculated by unpaired t test. Line represents mean.

To confirm that the PVRIG independent role for PVRL2 in promoting tumor growth is through the regulation of immune cells, we performed an *in vitro* NK cell killing assays on the TRAMP-C2 WT vs. *Pvrl2* KO cells. WT and *Pvrig* KO NK cells were mixed 1:1 with WT and *Pvrl2* KO TRAMP-C2 cells (Fig. 5J). PVRL2 loss in the TRAMP-C2 significantly enhanced killing by both WT and *Pvrig* KO NK cells (Fig. 5K). In contrast, the loss of PVRIG had no significant impact on the NK killing of WT TRAMP-C2 cells (Fig. 5K). Therefore, PVRL2’s PVRIG independent roles are through the direct regulation of immune cells, at least in the case of NK cells.

### PVRL2 loss and TIGIT blockade function cooperatively to inhibit tumor growth

Some reports have suggested that PVRL2 can also bind and regulate TIGIT (17, 18). Therefore, we asked if PVRL2’s PVRIG independent function could be through TIGIT. To do so, we injected WT MC38 cells into WT C57BL/6 mice followed by multiple injections of anti-TIGIT blocking antibody (Fig. 6A). Anti-TIGIT led to a significant reduction in tumor growth and extension in survival (Fig. 6B-D). Next, we repeated the experiments in the *Pvrig* KO background to determine if there was any collaborative effect of PVRIG with TIGIT underlying PVRL2’s immune suppressive role. With combined PVRIG loss and TIGIT blockade, *Pvrl2* KO still significantly retarded tumor growth and extended survival showing that PVRL2 can function independently of both PVRIG and TIGIT (Fig. 6E-G). Further, combination of *Pvrig* KO plus anti-TIGIT blockade did not show any further inhibition of tumor growth or extension of survival relative to anti-TIGIT alone (Supplementary Fig. S4G). However, the combination of PVRL2 loss in the tumor cells with anti-TIGIT blockade showed a significantly greater effect than either alone (Fig. 6E-G). These data suggest that PVRL2 is not acting through TIGIT, but instead in a parallel pathway with both playing important roles in suppressing the anti-tumor immune response. In contrast, PVRIG appears to play a relatively minor role.

**Figure 6:**
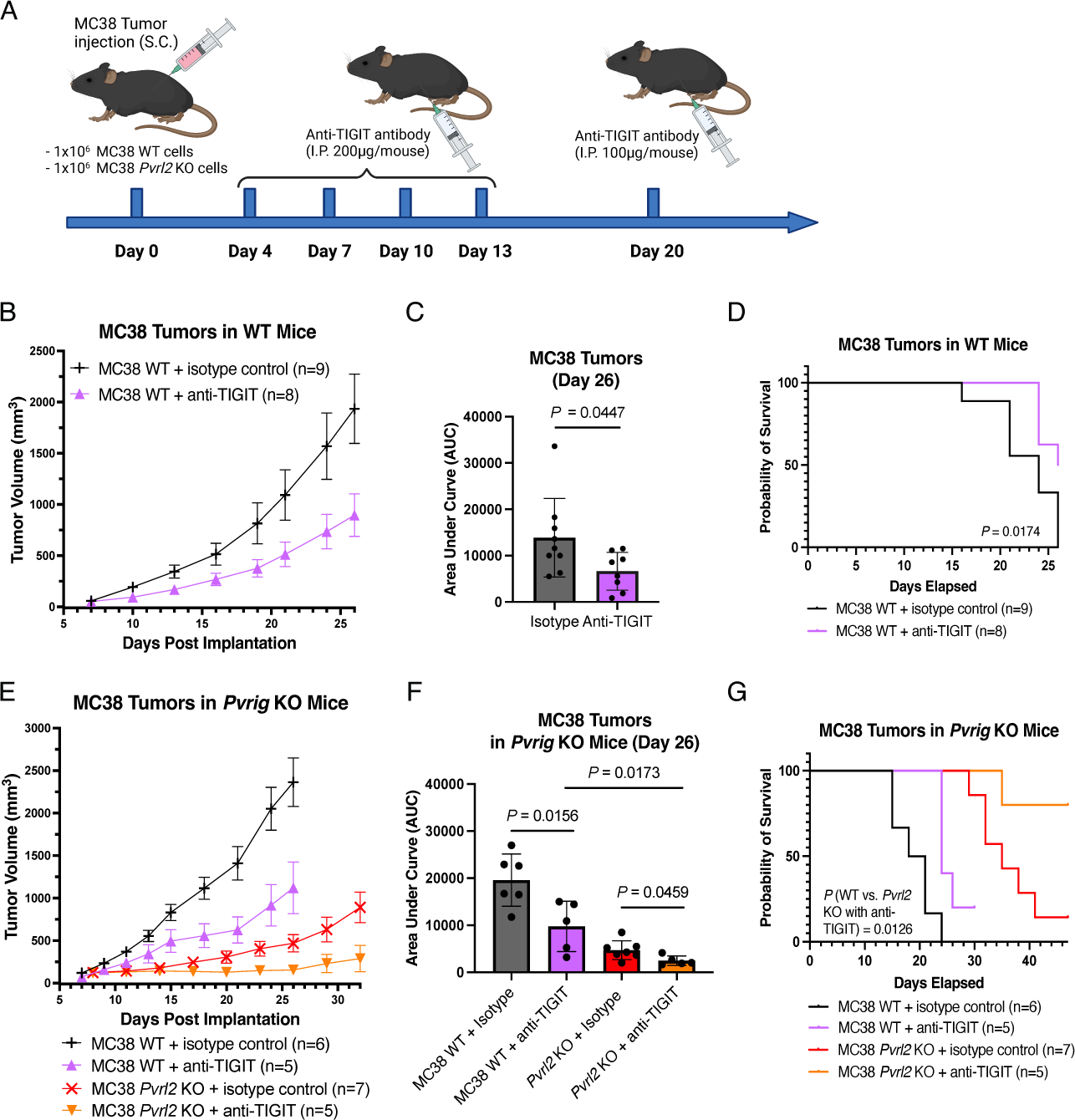
PVRL2 loss and TIGIT blockade function cooperatively to inhibit tumor growth. **(A)** Schematic of experiment design in (B-G): After subcutaneous injection of 1 × 10^6^ MC38 WT and *Pvrl2* KO cells in WT C57BL/6 or *Pvrig* KO mice, starting on day 4, mice were treated with serial doses of anti-TIGIT blocking antibody or isotype control at the indicated doses and schedule. **(B)** Average tumor volume over time following subcutaneous injection of 1 × 10^6^ MC38 WT cells with and without anti-TIGIT treatment in WT C57BL/6 mice. Error bars represent SEM. **(C)** Area under the curves of the MC38 tumors from (B) on day 26. Dots represent individual mice. *P* value is calculated by unpaired t test. Error bars represent SD. **(D)** Mouse survival curves following injections as described in (B). *P* value is calculated by log rank test. **(E)** Average tumor volume over time following subcutaneous injection of 1 × 10^6^ MC38 WT and *Pvrl2* KO cells with and without anti-TIGIT treatment in *Pvrig* KO mice. Error bars represent SEM. **(F)** Area under the curves of the MC38 tumors from (E) on day 26. Dots represent individual mice. *P* values are calculated by unpaired t test. Error bars represent SD. **(G)** Mouse survival curves following injections as described in (E). *P* values are calculated by log rank test.

Given that antibody blockade can be incomplete, we further validated any potential role for TIGIT by using CRISPR mutagenesis to knock out the *Tigit* gene in the *Pvrig* KO NK cells, and repeated *in vitro* NK cell killing assays on the TRAMP-C2 WT and *Pvrl2* KO cells (Supplementary Fig. S4H). Even in the absence of both PVRIG and TIGIT, PVRL2 loss showed an equally strong suppression of NK cell driven killing (Supplementary Fig. S4I and S4J). CD96 has also been suggested as a potential receptor for Nectin family proteins, although its immunoregulatory role has been controversial (21). Therefore, we also knocked out *Cd96* in the *Pvrig* KO NK cells and performed the NK cell killing assay (Supplementary Fig. S4H). Like TIGIT, the loss of CD96 together with PVRIG did not impact PVRL2’s ability to suppress NK cell killing (Supplementary Fig. S4I and S4J). Together, these data show that PVRL2 is acting to suppress antitumor immunity through mechanisms that are independent of the known Nectin family receptors on immune cells including PVRIG, TIGIT, and CD96.

### Combined loss of PVRL2 and PVR does not further inhibit tumor growth

The PVRL2 paralog, PVR is believed to be the primary ligand of TIGIT (17, 18). Flow cytometry confirmed that the loss of PVRL2 in MC38 cells doesn’t have an impact on PVR expression both *in vitro* and *in vivo* (Supplementary Fig. S5A-S5D). To test the impact of PVR loss alone and in combination with PVRL2 loss and/or TIGIT blockade, we used CRISPR/Cas9 mutagenesis to target the *Pvr* gene in MC38 cells. PVR loss was confirmed by flow cytometry (Supplementary Fig. S5A). The loss of PVR reduced tumor growth and extended survival, but to a lesser degree than PVRL2 (Fig. 7A-C). This effect was through the immune system, as there was no difference in growth between WT and *Pvr* KO tumor cells in the immunodeficient NCG mouse background (Supplementary Fig. S5E and S5F). However, PVR appeared to have a similar effect on growth and survival in the *Rag1* KO versus WT mouse background, suggesting that PVR is functioning largely through the innate rather than adaptive immune response (Fig. 7D-F). Combining PVR loss with anti-TIGIT blockade showed no additional inhibition of tumor growth or extension of survival, demonstrating that all PVR activity in regulating tumor growth is through TIGIT and vice versa (Fig. 7G-I). Combination of PVR and PVRIG loss also showed no additional effect relative to PVR loss alone (Supplementary Fig. S5I-S5K). Next, we tested the combination of knocking out *Pvrl2* and *Pvr* in the same tumor cells, expecting it to phenocopy the combination of PVRL2 loss with anti-TIGIT blockade. Surprisingly though, there was no additive effect in tumor growth or survival (Fig. 7A-C). If anything, there was a slight increase in tumor growth in the double KO relative to *Pvrl2* KO alone, although this did not show in survival or in *Rag1* KO background (Fig. 7A-F). All the cells produced tumors at a similar rate in NCG mice, confirming activity through targeting the anti-tumor immune response (Supplementary Fig. S5G and S5H). These data show that although PVR functions through TIGIT, combined loss of PVRL2 and PVR does not have the same positive impact as loss of PVRL2 combined with TIGIT blockade.

**Figure 7:**
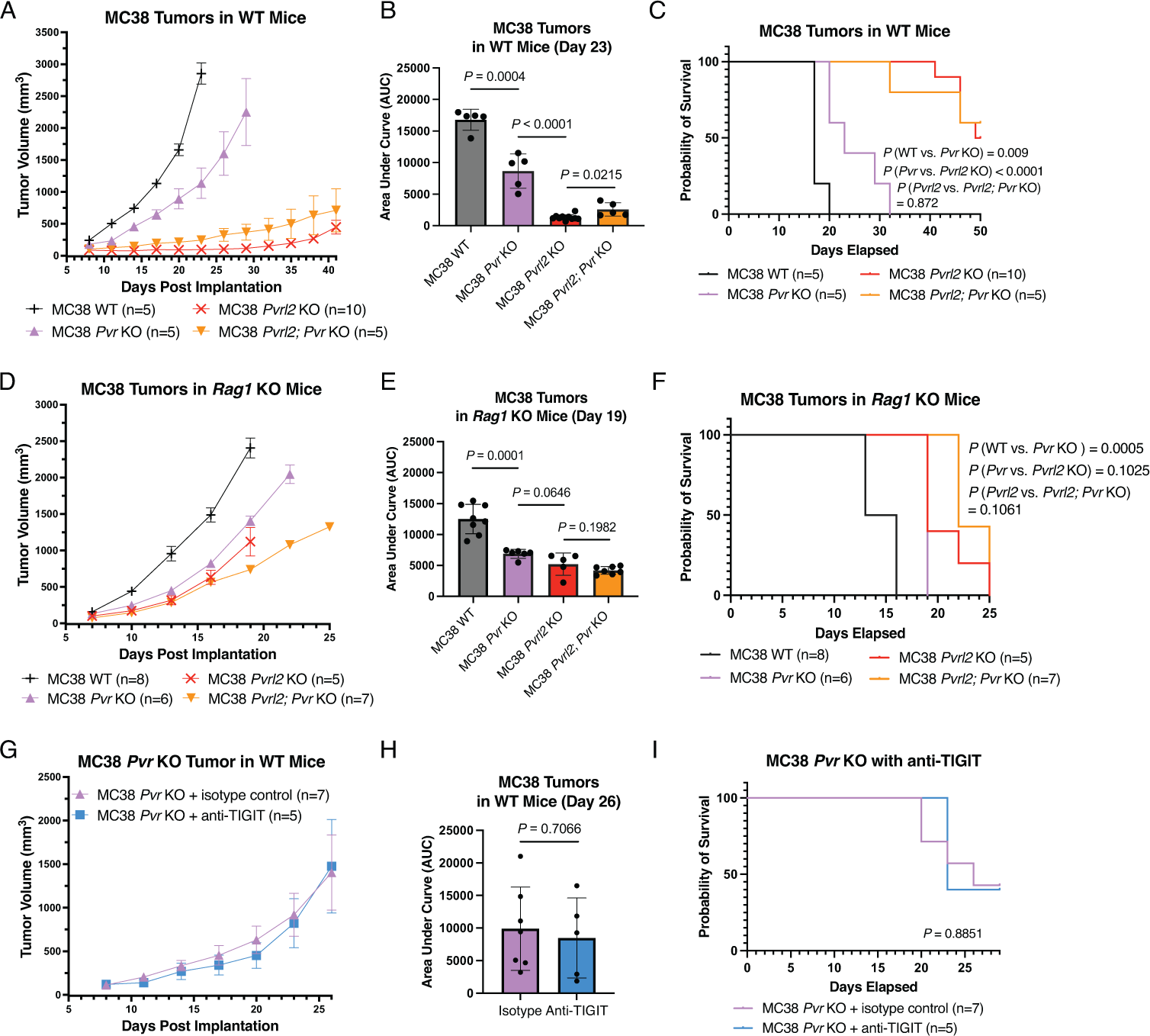
Combined loss of PVRL2 and PVR loss does not further inhibit tumor growth. **(A)** Average tumor volume over time following subcutaneous injection of 1 × 10^6^ MC38 WT, *Pvrl2* KO, *Pvr* KO and *Pvrl2*; *Pvr* KO cells in WT C57BL/6 mice. Error bars represent SEM. **(B)** Area under the curves of the MC38 tumors from (A) on day 23. Dots represent individual mice. *P* values are calculated by unpaired t test. Error bars represent SD. **(C)** Mouse survival curves following injections as described in (A). *P* values are calculated by log rank test. **(D)** Average tumor volume over time following subcutaneous injection of 1 × 10^6^ MC38 WT, *Pvrl2* KO, *Pvr* KO and *Pvrl2*; *Pvr* KO cells in *Rag1* KO mice. Error bars represent SEM. **(E)** Area under the curves of the MC38 tumors from (D) on day 19. Dots represent individual mice. *P* values are calculated by unpaired t test. Error bars represent SD. **(F)** Mouse survival curves following injections as described in (D). *P* values are calculated by log rank test. **(G)** Average tumor volume over time following subcutaneous injection of 1 × 10^6^ MC38 *Pvr* KO cells with and without anti-TIGIT treatment in WT C57BL/6 mice. Error bars represent SEM. **(H)** Area under the curves of the tumors from (G) on day 26. Dots represent individual mice. *P* value is calculated by unpaired t test. Error bars represent SD. **(I)** Mouse survival curves following injections as described in (G). *P* value is calculated by log rank test.

## Discussion

Our results uncover a profound impact of PVRL2 on suppressing the anti-tumor immune response across multiple tumor models that is largely independent of its known receptor, PVRIG. PVRL2 was initially identified in the late 1990s as an adhesion molecule belonging to the Nectin and Nectin-like family that supports cell-cell-junction formation (13, 52). In recent years, accumulating evidence has pointed toward a role for PVRL2 in cancer and the modulation of anti-tumor immunity (14, 18, 25, 44). Elevated levels of PVRL2 are found in many cancer types, including acute myeloid leukemia, multiple myeloma, and epithelial cancers such as colorectal cancer, melanoma, lung cancer, endometrial cancer, breast cancer, prostate cancer, and ovarian cancer (25, 53-56). While PVRL2’s function in tumor development has been attributed to its ability to bind the co-inhibitory receptor PVRIG on immune cells, there has been little evidence to support that conclusion *in vivo*. Previous research has been limited to *in vitro* experiments showing that anti-PVRL2 blockade can stimulate the activation of CD8 T cells when co-cultured with the human melanoma Mel-624 cell line (25), or enhance PBMC-mediated lysis of hepatocellular carcinoma (HCC) cell lines (57). To our knowledge, this study represents the first direct investigation into the mechanism of PVRL2 *in vivo* using *Pvrl2* KO mouse tumor models, in turn providing compelling evidence for its potential as a therapeutic target.

Tumor-derived exosomes have emerged as an important mechanism by which tumor cells suppress anti-tumor immunity by carrying immunosuppressive molecules, such as PD-L1 (5, 6, 9, 10), and their expression on exosomes have been identified as potential biomarkers for responses to ICIs in patients (58). In this study, we identify PVRL2 as another immune suppressive molecule on exosomes released by multiple human and mouse tumor cell lines, as well as primary prostate tumor tissue. Our *in vivo* experiments reveal that exosomal PVRL2 can significantly promote tumor growth, although the contribution is relatively small compared to the substantial impact observed for PVRL2 on cells. This result indicates that the role of PVRL2 in promoting tumor growth primarily occurs though cell surface PVRL2 rather than via exosomes, which led us to focus on elucidating the mechanisms underlying the immunosuppressive role of cellular PVRL2 in tumor development.

By using multiple syngeneic mouse tumor models, we demonstrate that PVRL2 significantly promotes tumor growth and suppresses the anti-tumor immune response. Specifically, PVRL2 exerts its tumor-promoting effects by suppressing CD8 T cells and NK cells. However, surprisingly, our findings indicate PVRL2’s function is predominantly independent of PVRIG, as evidenced by the minimal impact of PVRIG loss on the *Pvrl2* knockout phenotype. Previous literatures have reported slower tumor growth in *Pvrig* KO mice for both MC38 and B16F10 tumors, although the difference was quite small (24, 50, 51). Our experiments similarly reveal slightly reduced growth of B16F10 WT tumors in *Pvrig* KO mice. However, we do not observe any impact on the growth of MC38 and TRAMP-C2 tumors with the loss of PVRIG. One potential explanation for this discrepancy is the difference in the number of tumor cells injected per mouse in our study (1 million cells) compared to the previous literatures (0.2 or 0.5 million cells). The higher tumor burden may overwhelm the suppressive properties of PVRIG, resulting in even less noticeable differences in tumor growth. Importantly, even in the absence of PVRIG, *Pvrl2* KO tumors grew much slower than their WT counterparts in all three models, even at the high tumor cell doses. Therefore, there must be additional receptor(s) for PVRL2. An obvious candidate is TIGIT given previous work suggesting low affinity binding as well as upregulation of TIGIT on immune cells, particularly CD8 T cells, upon PVRIG blockade in co-culture experiments with tumor cells (25). However, TIGIT blockade alone or in combination with PVRIG loss failed to inhibit the tumor promoting impact of PVRL2. Thus, further studies will be required to uncover the primary receptor(s) that mediates PVRL2 function.

The primary ligand for TIGIT is the PVRL2 paralog PVR (17, 18). Indeed, we find that PVR function is lost with blockade of TIGIT and vice-versa. Therefore, PVR’s ability to suppress the immune response appears to be entirely through TIGIT, in stark contrast to the PVRL2-PVRIG axis. Notably, the impact of PVRL2 loss on tumor growth was significantly greater than that of PVR loss. Furthermore, consistent with prior literature (59), we show that PVR appears to act predominantly through the innate immune response, as its impact on tumor growth was unaffected in the *Rag1* KO background, where there is no adaptive immune response. In contrast, we show that PVRL2 affects both the adaptive and innate immune responses presumably through direct regulation of CD8 T and NK cells, although discovery of the PVRL2 receptor(s) will be required to confirm that.

We found that the most profound suppressive effect on tumor growth was produced by a combination of PVRL2 loss and TIGIT blockade. However, this effect was not recapitulated with the combination of PVRL2 and PVR loss. If anything, the double KO led to slightly increased growth compared to PVRL2 loss alone. The most likely explanation for this seemingly contradictory finding is a role for the co-stimulatory receptor, DNAM-1 in transmitting an immune-promoting signal from PVRL2 and PVR (14–16). It is possible the loss of either PVRL2 or PVR alone has little negative consequence on DNAM-1 activation, but the loss of both leads to loss of the immune-activating signal. In either case, the immune inhibitory functions of PVR and PVRL2 seem to be much greater than their immune promoting functions given that the double KO tumors still grow much slower than WT tumors.

Our preclinical results in mouse models provide strong evidence for the therapeutic potential of targeting PVRL2 to reactivate the anti-tumor immune response. Our data show that this potential therapeutic impact of PVRL2 inhibition can be further enhanced by TIGIT, but not PVR, blockade. Thus, our data provide strong rationale for combinatorial PVRL2 and TIGIT inhibition. In future studies, it will be important to determine how such dual inhibition will interact with anti-PD-1 or PD-L1 blockade. Indeed, prior results have suggested cooperation between anti-TIGIT, anti-PVRIG, and anti-PD-1. Our data would suggest a substantially greater impact of anti-TIGIT, anti-PVRL2, and anti-PD-1.

## Authors’ Contributions

J.Y. and R.B. conceived the project and designed the experiments. J.Y. performed the exosome preparation from cells, gene enrichment analysis for mass spectrometry, and all biochemical, cellular experiments, animal experiments, and their data analysis. J.R.B. and L.L.K. performed the mass spectrometry and analysis. L.W. performed the IF sample preparation, staining, imaging, and image analysis, and nucleofection in NK cells for generating *Tigit* and *Cd96* knockout. H.D. contributed to the exosome injection and generating MC38 *Pdl1* KO cells. C.D.B. prepared exosome samples from primary tumor tissues. L.F. provided helpful advice and guidance throughout the project. O.A.A. and L.L.L. advised the NK cytotoxicity assay. J.Y. and R.B. wrote the manuscript. J.R.B., L.W., H.D. and C.D.B. provided detailed methods for the experiments they performed. All authors proofread and provided feedback.

## Acknowledgments

We thank all members of the UCSF Tumor Immunology Joint Meeting for helpful discussions and suggestions. We thank Carolyn Sangokoya, Ryan Boileau, Deniz Goekbuget for their helpful advice and feedback throughout the project. We thank Dr. Alexander Marson for kindly sharing the *Rag1* KO mice. We thank Dr. Jeffrey Schlom for kindly providing the MC38 cell line. We thank UCSF BIOS tissue bank for sharing the primary prostate tumor tissues. The project is funded by NIH U01 grant (U01CA244452).

