## Supplemental Figures and Table for "PVRL2 Suppresses Anti-tumor Immunity Through PVRIG- and TIGIT-Independent Pathways"

### Supplementary Information

A

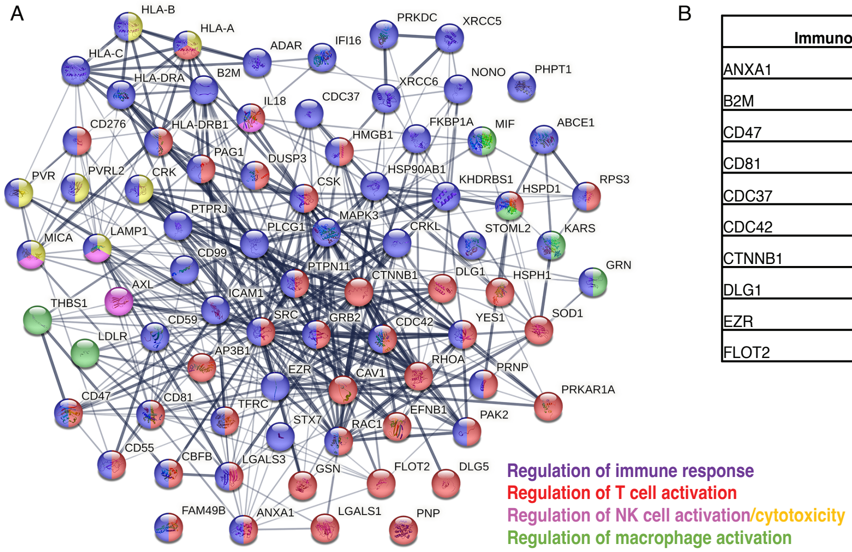


B

| Immunoregulators Shared by All Samples (28) | | |
|---|---|---|
| ANXA1 | GSN | PRKAR1A |
| B2M | HSP90AB1 | PVRL2 |
| CD47 | HSPH1 | RPS3 |
| CD81 | LGALS1 | SOD1 |
| CDC37 | LGALS3 | SRC |
| CDC42 | MAPK3 | STX7 |
| CTNNB1 | MIF | TFRC |
| DLG1 | NONO | YES1 |
| EZR | PHPT1 | |
| FLOT2 | PNP | |

C

| | PC3 | | SK-MEL-28 | |
|---|---|---|---|---|
| **Gene Names** | -Lg (*P* value) | $Log_2$ (Fold Change) | -Lg (*P* value) | $Log_2$ (Fold Change) |
| DLG1 | 3.374 | 4.238 | 4.044 | 5.291 |
| YES1 | 4.215 | 3.666 | 3.002 | 3.850 |
| PVRL2 | 1.752 | 3.354 | 3.118 | 2.601 |
| CTNNB1 | 3.193 | 2.823 | 3.941 | 3.163 |

D

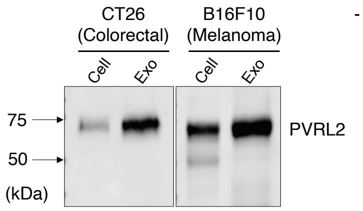


**Supplementary Figure S1:** Proteomic analysis identifies PVRL2 on tumor-derived exosomes.

**(A)** Functional association network of the 78 proteins identified on PC3 and SK-MEL-28 exosomes that have direct immunoregulatory function, as identified by gene enrichment analysis (by ShinyGo 0.77), shown as indicated functional categories (generated from String-db). The line thickness represents the strength of data support for functional and physical associations. **(B)** List of the 28 immunoregulatory proteins shared by exosomes from PC3, SK-MEL-28, TRAMP-C2 and primary tumor slices. **(C)** 4 immunoregulatory proteins over 2-fold enriched ($Log_2$ (Fold Change) > 1) in both PC3 and SK-MEL-28 exosomes relative to cells. Fold change and *P* values are indicated. **(D)** Western blot analysis for PVRL2 in the cells and exosomes (exo) from CT26 and B16F10 cell lines. 40 µg of total protein was loaded for each sample.

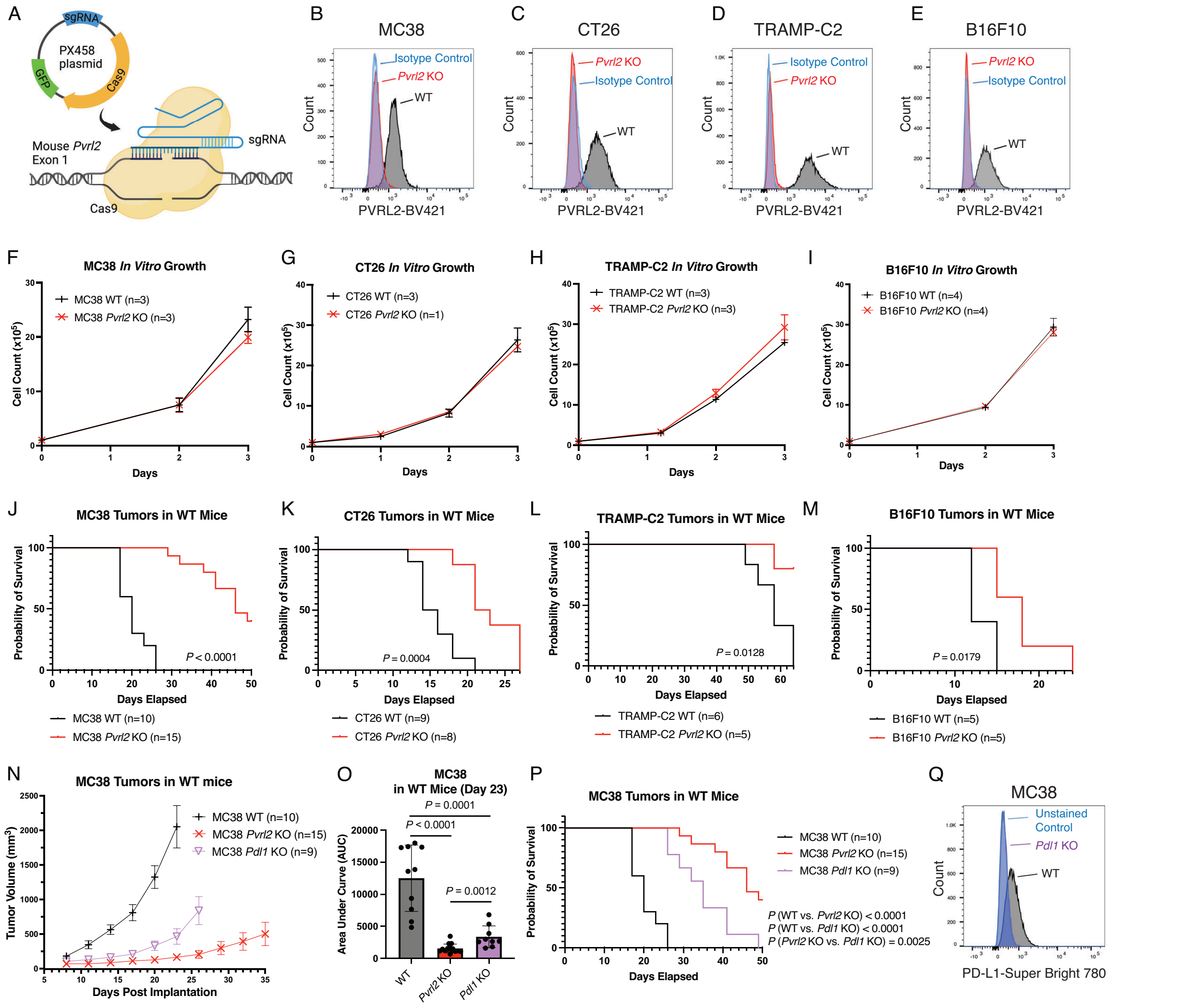


**Supplementary Figure S2:** PVRL2 promotes tumor growth through an immune-dependent mechanism. (See next page for caption.)

**Supplementary Figure S2:** PVRL2 promotes tumor growth through an immune-dependent mechanism.

**(A)** Schematic of generating *Pvrl2* KO cells by CRISPR/Cas9 gene editing by using PX458 plasmid expressing sgRNA targeting *Pvrl2* exon 1. **(B-E)** Flow cytometry of PVRL2 surface expression on MC38 (B), CT26 (C), TRAMP-C2 (D) and B16F10 (E) WT and *Pvrl2* KO cells. **(F-I)** *In vitro* cell growth of MC38 (F), CT26 (G), TRAMP-C2 (H) and B16F10 (I) WT and *Pvrl2* KO cells as shown by cell counts over days. **(J-M)** Mouse survival curves following subcutaneous injections of $1 \times 10^6$ MC38 (J), TRAMP-C2 (L), and B16F10 (M) WT and *Pvrl2* KO cells in WT C57BL/6 mice, and CT26 (K) WT and *Pvrl2* KO cells in WT BALB/cJ mice. *P* values are calculated by log rank test. **(N)** Average tumor volume over time following subcutaneous injection of $1 \times 10^6$ MC38 WT, *Pvrl2* KO, and *Pdl1* KO cells in WT C57BL/6 mice. Error bars represent SEM. **(O)** Area under the curves of the MC38 tumors from (N) calculated on day 23. Dots represent individual mice. *P* values are calculated by unpaired t test. Error bars represent SD. **(P)** Mouse survival curves following injections as described in (N). *P* values are calculated by log rank test. **(Q)** Flow cytometry of PD-L1 surface expression on MC38 WT and *Pdl1* KO cells.

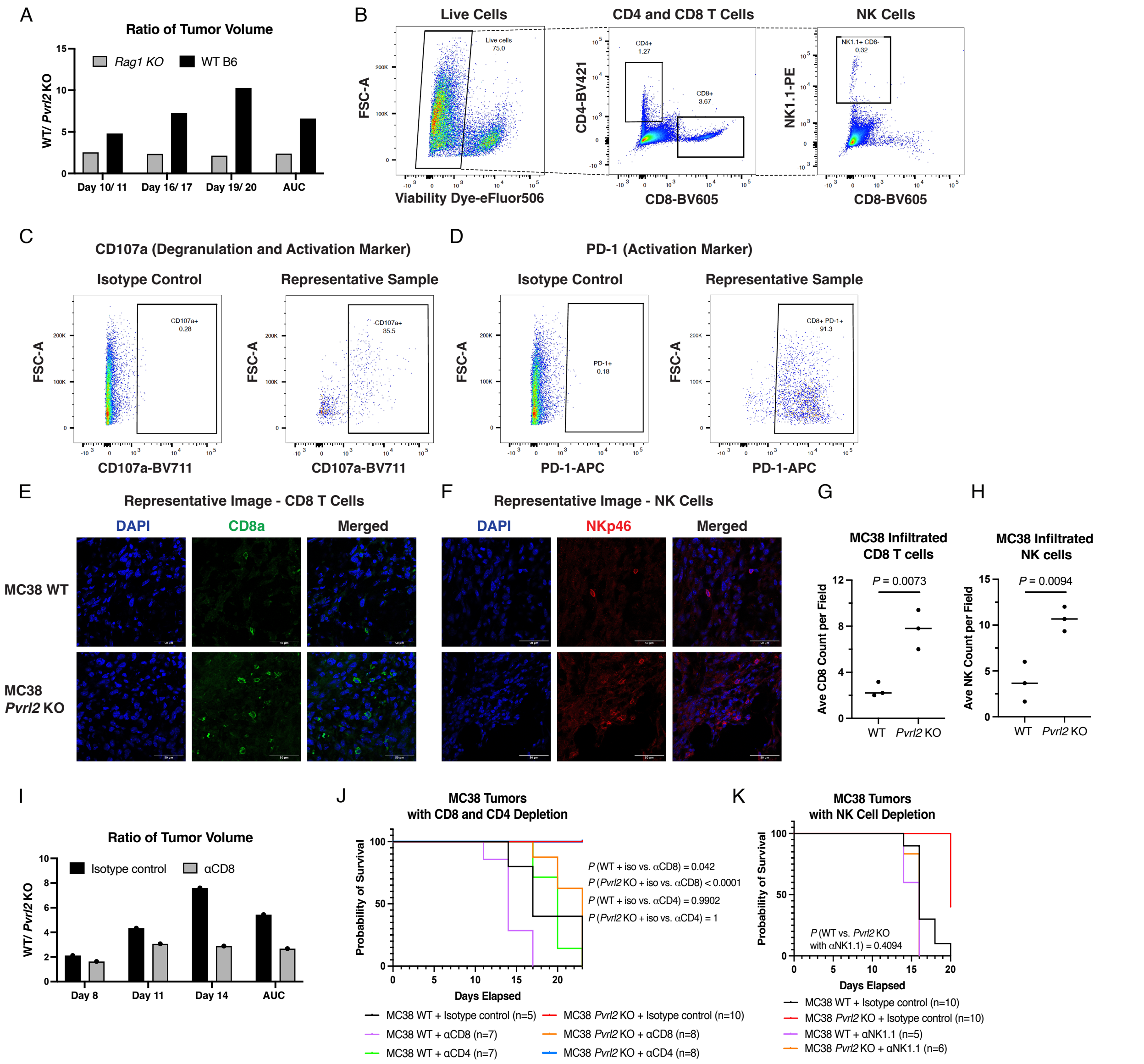


**Supplementary Figure S3:** PVRL2 regulates CD8 T cell and NK cell activation. (See next page for caption.)

**Supplementary Figure S3:** PVRL2 regulates CD8 T cell and NK cell activation.

**(A)** Ratios of the average WT/ *Pvrl2* KO tumor volumes on day 10, 16, and 19 from Fig. 4 (A) and day 11, 17, and 20 from Fig. 2 (A), and ratios of the AUC of WT/ *Pvrl2* KO tumors on day 19 from Fig. 4 (B) and day 20 from Fig. 2 (B). **(B)** Gating strategy for flow cytometry analysis of tumor infiltrated CD4, CD8 T cells, and NK cells. **(C-D)** Representative flow plots of T cells and NK cells stained for CD107a (C) and PD-1 (D) over isotype-matched antibody controls. **(E-F)** Representative immunofluorescence (IF) images for CD8 T cells (E) and NK cells (F) in MC38 WT and *Pvrl2* KO tumors collected on day 25 or 27. Scale bar = 50 μm. **(G-H)** IF quantification of the average numbers of CD8 T cells (G) and NK cells (H) in 3 imaged fields of MC38 WT and *Pvrl2* KO tumors (n=3). *P* value is calculated by unpaired t test. Error bars represent SD. Line represents mean. **(I)** Ratios of the average WT/ *Pvrl2* KO tumor volumes at the indicate timepoints from Fig. 4 (K) and ratios of the AUC of WT/ *Pvrl2* KO tumors on day 17 from Fig. 4 (M). **(J)** Mouse survival curves following injections as described in Fig. 4 (K,L). *P* values are calculated by log rank test. **(K)** Mouse survival curves following injections as described in Fig. 4 (N). *P* values are calculated by log rank test.

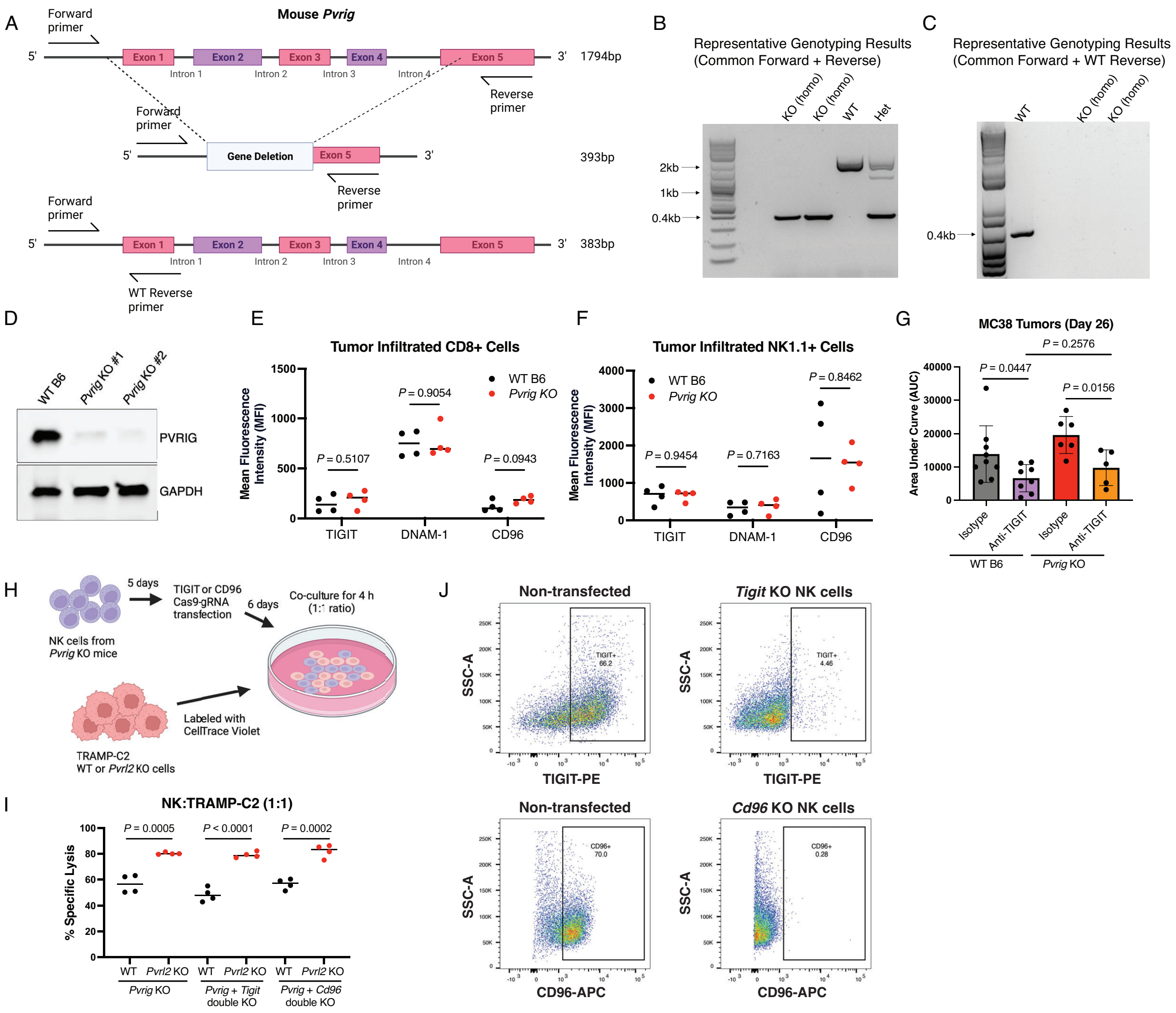


**Supplementary Figure S4:** PVRL2 functions independently of PVRIG and TIGIT pathways. (See next page for caption.)

**Supplementary Figure S4:** PVRL2 functions independently of PVRIG and TIGIT pathways.

**(A)** Schematic of the deletion of *Pvrig* in C57BL/6 mice and genotyping strategy by PCR. **(B)** Representative image of the genotyping results using the combination of common forward and common reverse primers, showing two homozygous *Pvrig* KO, one WT and one heterozygous KO samples. **(C)** Representative image of the genotyping results using the combination of common forward and WT specific reverse primers, showing one WT and two homozygous *Pvrig* KO samples. **(D)** Western blot analysis for PVRIG protein in the splenocytes from one WT and two homozygous *Pvrig* KO mice. 30 μg of total protein was loaded for each sample. GAPDH was used as the loading control. **(E-F)** Flow cytometric analysis of the Mean Fluorescence Intensity (MFI) of TIGIT, DNAM-1, and CD96 expression on the CD8+ (E) and NK1.1+ (F) sub-populations, in the CD45+ cells isolated from MC38 WT tumors in WT mice (n = 4) and *Pvrig* KO mice (n = 4). *P* values are calculated by unpaired t test. Line represents mean. **(G)** Area under curves of MC38 WT tumors with anti-TIGIT antibody treatment or isotype-matched control antibody in WT and *Pvrig* KO mice in Fig. 6 (B,E) on day 26. *P* values are calculated by unpaired t test. Error bars represent SD. **(H)** Schematic of experiment design in (I): *Pvrig* KO NK cells isolated from *Pvrig* KO mice were transfected with *Tigit* or *Cd96* gRNAs for gene CRISPR knockout on day 5. 6 days after transfection, NK cells were co-cultured at 1:1 with TRAMP WT or *Pvrl2* KO tumor cells for 4 hours. **(I)** Percentage lysis of TRAMP-C2 WT and *Pvrl2* KO cells after co-culturing with *Pvrig* KO, *Pvrig*; *Tigit* double KO, or *Pvrig*; *Cd96* double KO NK cells at 1:1 ratio. Dots represent individual replicates. *P* values are calculated by unpaired t test. Line represents mean. **(J)** Validation of the TIGIT and CD96 KO efficiency by flow cytometry with TIGIT and CD96 staining on non-transfected NK cells compared to transfected NK cells 6 days post-transfection.

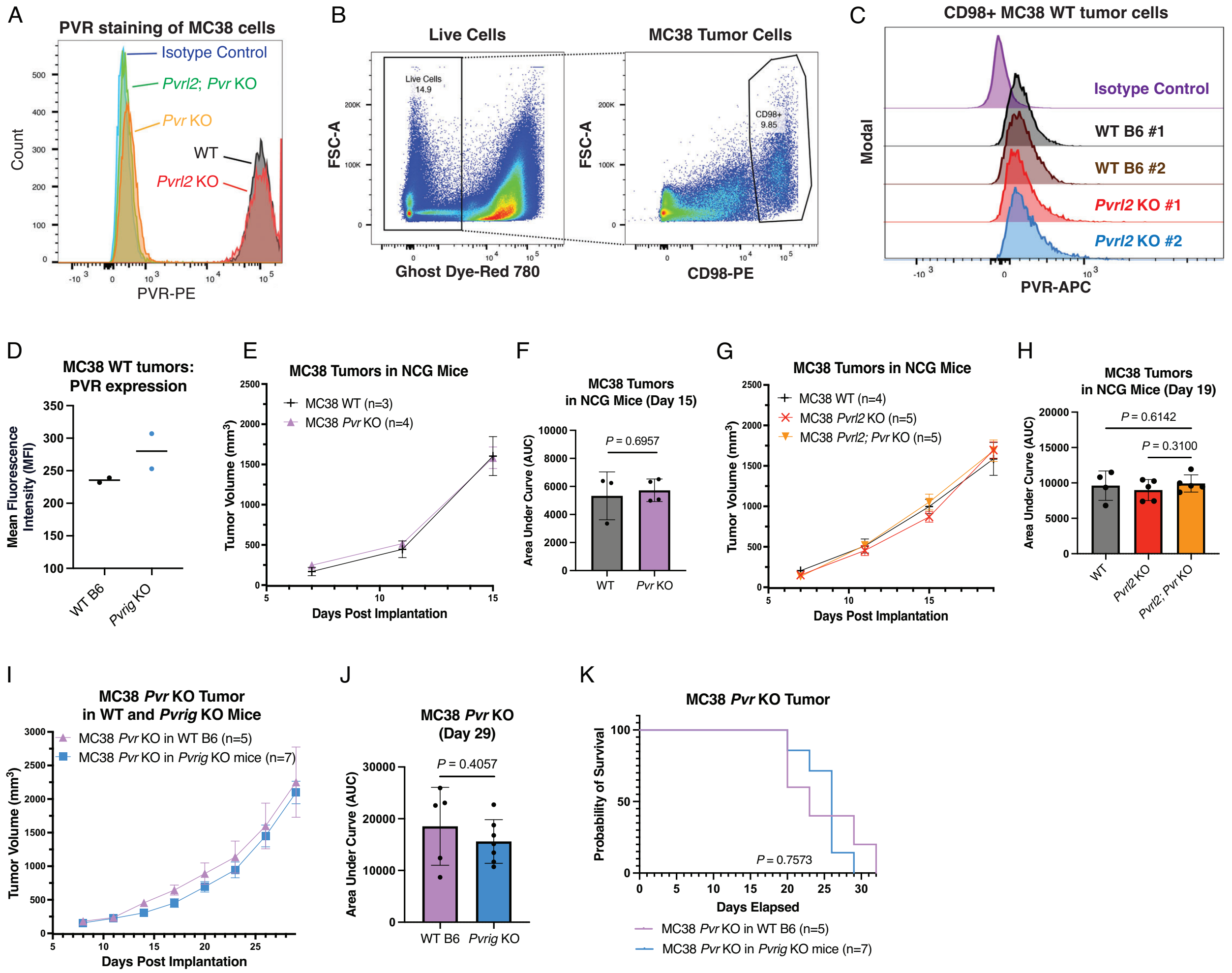


**Supplementary Figure S5:** Combined loss of PVRL2 and PVR loss does not further inhibit tumor growth. (See next page for caption.)

**Supplementary Figure S5:** Combined loss of PVRL2 and PVR loss does not further inhibit tumor growth.

**(A)** Flow cytometry with PVR surface staining on MC38 WT, *Pvrl2* KO, *Pvr* KO and *Pvrl2*; *Pvr* KO cell lines. **(B)** Gating strategy for flow cytometry analysis of MC38 tumor cells in (C). **(C)** Histogram flow plots of PVR expression levels on the CD98+ sub-population from MC38 WT and *Pvrl2* KO tumors in WT mice. **(D)** Flow cytometric analysis of the MFI of PVR expression on MC38 WT and *Pvrl2* KO tumors from (C). Line represents mean. **(E)** Average tumor volume over time following subcutaneous injection of $1 \times 10^6$ MC38 WT and *Pvr* KO cells in NCG mice. Error bars represent SEM. **(F)** Area under the curves of the tumors from (E) on day 15. *P* value is calculated by unpaired t test. Error bars represent SD. **(G)** Average tumor volume over time following subcutaneous injection of $1 \times 10^6$ MC38 WT, *Pvrl2* KO, and *Pvrl2*; *Pvr* KO cells in NCG mice. Error bars represent SEM. **(H)** Area under the curves of the tumors from (G) on day 19. *P* values are calculated by unpaired t test. Error bars represent SD. **(I)** Average tumor volume over time following subcutaneous injection of $1 \times 10^6$ MC38 *Pvr* KO cells in WT C57BL/6 mice and *Pvrig* KO mice. Error bars represent SEM. **(J)** Area under the curves of the MC38 *Pvr* KO tumors from (I) on day 29. *P* value is calculated by unpaired t test. Error bars represent SD. **(K)** Mouse survival curves following injections as described in (I). *P* value is calculated by log rank test.

**Supplementary Table S1**: Statistical analysis between groups of areas under curves (AUC) of tumor growth data (unpaired student's T test).

| Figures | Comparison | | *P* values |
|---|---|---|---|
| **Fig. 4M** | MC38 WT + isotype control | MC38 *Pvrl2* KO + isotype control | < 0.0001 |
| | MC38 WT + isotype control | MC38 WT + αCD8 | 0.0202 |
| | MC38 WT + isotype control | MC38 WT + αCD4 | 0.464 |
| | MC38 *Pvrl2* KO + isotype control | MC38 *Pvrl2* KO + αCD8 | < 0.0001 |
| | MC38 *Pvrl2* KO + isotype control | MC38 *Pvrl2* KO + αCD4 | 0.0099 |
| | MC38 WT + αCD8 | MC38 *Pvrl2* KO + αCD8 | 0.0001 |
| | MC38 WT + αCD4 | MC38 *Pvrl2* KO + αCD4 | < 0.0001 |
| **Fig. 4O** | MC38 WT + isotype control | MC38 *Pvrl2* KO + isotype control | 0.0003 |
| | MC38 WT + isotype control | MC38 WT + αNK1.1 | < 0.0001 |
| | MC38 WT + isotype control | MC38 *Pvrl2* KO + αNK1.1 | < 0.0001 |
| | MC38 *Pvrl2* KO + isotype control | MC38 WT + αNK1.1 | < 0.0001 |
| | MC38 *Pvrl2* KO + isotype control | MC38 *Pvrl2* KO + αNK1.1 | < 0.0001 |
| | MC38 WT + αNK1.1 | MC38 *Pvrl2* KO + αNK1.1 | 0.1413 |
| **Fig. 5B** | MC38 WT tumor in WT mice | MC38 *Pvrl2* KO tumor in WT mice | 0.0063 |
| | MC38 WT tumor in WT mice | MC38 WT tumor in *Pvrig* KO mice | 0.8742 |
| | MC38 WT tumor in WT mice | MC38 *Pvrl2* KO tumor in *Pvrig* KO mice | 0.0235 |
| | MC38 *Pvrl2* KO tumor in WT mice | MC38 WT tumor in *Pvrig* KO mice | 0.0003 |
| | MC38 *Pvrl2* KO tumor in WT mice | MC38 *Pvrl2* KO tumor in *Pvrig* KO mice | 0.1378 |
| | MC38 WT tumor in *Pvrig* KO mice | MC38 *Pvrl2* KO tumor in *Pvrig* KO mice | 0.0023 |
| **Fig. 5E** | TRAMP-C2 WT tumor in WT mice | TRAMP-C2 *Pvrl2* KO tumor in WT mice | 0.0018 |
| | TRAMP-C2 WT tumor in WT mice | TRAMP-C2 WT tumor in *Pvrig* KO mice | 0.4866 |
| | TRAMP-C2 WT tumor in WT mice | TRAMP-C2 *Pvrl2* KO tumor in *Pvrig* KO mice | 0.0048 |
| | TRAMP-C2 *Pvrl2* KO tumor in WT mice | TRAMP-C2 WT tumor in *Pvrig* KO mice | < 0.0001 |
| | TRAMP-C2 *Pvrl2* KO tumor in WT mice | TRAMP-C2 *Pvrl2* KO tumor in *Pvrig* KO mice | 0.3162 |
| | TRAMP-C2 WT tumor in *Pvrig* KO mice | TRAMP-C2 *Pvrl2* KO tumor in *Pvrig* KO mice | < 0.0001 |
| **Fig. 5H** | B16F10 WT tumor in WT mice | B16F10 *Pvrl2* KO tumor in WT mice | 0.0046 |
| | B16F10 WT tumor in WT mice | B16F10 WT tumor in *Pvrig* KO mice | 0.0436 |
| | B16F10 WT tumor in WT mice | B16F10 *Pvrl2* KO tumor in *Pvrig* KO mice | 0.0023 |
| | B16F10 *Pvrl2* KO tumor in WT mice | B16F10 WT tumor in *Pvrig* KO mice | 0.0309 |
| | B16F10 *Pvrl2* KO tumor in WT mice | B16F10 *Pvrl2* KO tumor in *Pvrig* KO mice | 0.3398 |

| | | | |
|---|---|---|---|
| | B16F10 WT tumor in *Pvrig* KO mice | B16F10 *Pvrl2* KO tumor in *Pvrig* KO mice | 0.0028 |
| **Fig. 6F** | MC38 WT + isotype control | MC38 WT + anti-TIGIT | 0.0156 |
| | MC38 WT + isotype control | MC38 *Pvrl2* KO + isotype control | < 0.0001 |
| | MC38 WT + isotype control | MC38 *Pvrl2* KO + anti-TIGIT | < 0.0001 |
| | MC38 WT + anti-TIGIT | MC38 *Pvrl2* KO + isotype control | 0.0434 |
| | MC38 WT + anti-TIGIT | MC38 *Pvrl2* KO + anti-TIGIT | 0.0173 |
| | MC38 *Pvrl2* KO + isotype control | MC38 *Pvrl2* KO + anti-TIGIT | 0.0459 |
| **Fig. 7B** | MC38 WT | MC38 *Pvr* KO | 0.0004 |
| | MC38 WT | MC38 *Pvrl2* KO | < 0.0001 |
| | MC38 WT | MC38 *Pvrl2; Pvr* KO | < 0.0001 |
| | MC38 *Pvr* KO | MC38 *Pvrl2* KO | < 0.0001 |
| | MC38 *Pvr* KO | MC38 *Pvrl2; Pvr* KO | 0.0016 |
| | MC38 *Pvrl2* KO | MC38 *Pvrl2; Pvr* KO | 0.0215 |
| **Fig. 7E** | MC38 WT | MC38 *Pvr* KO | 0.0001 |
| | MC38 WT | MC38 *Pvrl2* KO | 0.0001 |
| | MC38 WT | MC38 *Pvrl2; Pvr* KO | < 0.0001 |
| | MC38 *Pvr* KO | MC38 *Pvrl2* KO | 0.0646 |
| | MC38 *Pvr* KO | MC38 *Pvrl2; Pvr* KO | < 0.0001 |
| | MC38 *Pvrl2* KO | MC38 *Pvrl2; Pvr* KO | 0.1982 |
